# Clonal selection supported by single cell Dna sequencing reveals hormonal adaptation and resistance in locally advanced breast cancer During Neoadjuvant Aromatase Inhibition

**DOI:** 10.1101/2025.09.24.678208

**Authors:** Denise G. O’Mahony, Tom Lesluyes, Ina S. Brorson, Patrik Vernhoff, Ksenia Sokolova, Miriam Ragle Aure, Grethe G. Alnæs, Rebecca Mae Hoøen, Arvind Y.M. Sundaram, Chandra Theesfeld, Stephanie B. Geisler, Torill Sauer, Nazli Bahrami, Rahul Parulkar, Andliena Tahiri, Torben Lüders, Olga Troyanskaya, Charles Vaske, Peter van Loo, Jürgen Geisler, Vessela N. Kristensen

## Abstract

**Background:** The aromatase inhibitors (AI) letrozole and exemestane, are often used in sequence in targeting ER+ breast cancers. However resistance to AI poses a major barrier to sustained clinical benefit, while the biological mechanisms underlying the phenomenon remain largely unknown. In this study, we build on our clinical NeoLetExe trial, with the aim to investigate the molecular basis of resistance to AI, by analysing subclonal evolutionary dynamics during sequential treatment.

**Methods:** We use whole-exome sequencing (WES) data from 11 ER+ breast cancer patients and 3 timepoints of the Neoletexe trial to reconstruct cancer cell fraction–based subclonal composition. Single-cell DNA sequencing from matched tumour samples is used for validating identified clones and variants. Subclonal variants were annotated to genes by integrating evidence from public data and ExpectoSc. Pathway enrichment analysis using Human Base was conducted.

**Results:** Higher cancer cell fraction clone trajectories were significantly associated with reduced treatment response (p = 0.023). Clones reconstructed by WES were validated at 81% using single-cell DNA sequencing. Clones resistant to both letrozole and exemestane demonstrated PIK3CA/AKT/mTOR signaling activation, KRAS pathway dysregulation, hedgehog signaling, and androgen receptor pathways, alongside extensive immune activation and metabolic reprogramming. Drug-specific resistance patterns showed exemestane-resistant clones enriched for epigenetic control and miRNA-mediated silencing, while letrozole-resistant clones displayed metabolic dysregulation but notably lacked immune pathway activation. In contrast, treatment-sensitive clones maintained coordinated cell cycle control, preserved DNA damage responses, and retained immune signaling capacity. Analysis of FDA-approved breast cancer targets identified actionable alterations in *PIK3CA* (4 patients) and *AKT1* (1 patient) that persisted through AI treatment, with RNA expression analysis revealing 48 additional therapeutic targets spanning PI3K/AKT/mTOR, CDK4/6, DNA repair (BRCA1/2, ATM), and immune checkpoint pathways.

**Conclusion:** WES-based cancer cell fraction analysis successfully captured subclonal evolutionary trajectories during AI treatment, revealing drug-specific mechanisms and identifying key molecular players in endocrine therapy resistance. This work establishes a framework for precision oncology approaches by providing actionable therapeutic targets and advancing our understanding of resistance mechanisms to improve clinical outcomes in sequential AI therapy.

## Introduction

More than 70% of breast cancer cases are oestrogen receptor-positive (ER+), with tumour growth primarily driven by the steroid hormone oestradiol, the most potent form of free oestrogens in vivo^[1]^. However, ER+ breast cancer exhibits substantial biological heterogeneity, with wide variability in prognosis, histology, and treatment response^[1,2]^. In postmenopausal women with ER-positive and locally-advanced breast cancer, endocrine therapy for at least 6 months is used in the neoadjuvant setting to eliminate estrogen signalling, thereby suppressing tumour growth and clinical down-staging prior to surgery. Key endocrine agents include in general aromatase inhibitors (AIs), which function by inhibiting the aromatase enzyme (*CYP19*), thereby suppressing the peripheral conversion of androgens to oestrogen^[3]^. Among the commonly used AIs are letrozole (Femar®/Femara®) a non-steroidal competitive inhibitor and exemestane (Aromasin®), a steroidal irreversible inactivator of aromatase^[4]^. Letrozole competes reversibly with androgens for binding to aromatase, whereas exemestane forms a covalent bond with aromatase, leading to irreversible inactivation.

While AIs are effective in targeting ER+ breast cancers, resistance—whether intrinsic or acquired— poses a major barrier to sustained clinical benefit^[5]^. Mechanisms of resistance often involve the adaptive upregulation of pathways such as growth factor signalling and cross-talk with ER activity, reflecting the broader impact of oestrogen deprivation on cellular signalling networks^[6]^. In the neoadjuvant setting, distinct genetic alterations are associated with AI sensitivity and resistance^[7]^. Resistant tumours show characteristic patterns of proliferation, histological and intrinsic molecular subtypes, and enrichment of genes in TP53 signalling, DNA replication, and mismatch repair. Interestingly, although resistance can occur with any AI, some patients who relapse after treatment with non-steroidal AIs (e.g., letrozole or anastrozole) may still respond to a steroidal agent such as exemestane, and vice versa, thus, highlighting the potential clinical relevance of sequential AI use to extend treatment efficacy^[8,9]^. However, the biological mechanisms underlying this benefit remain poorly understood and may involve additional, as yet largely uncharacterised effects^[9–23]^.

Tumour subclones—genetically distinct cell populations within the same tumour—are key contributors to intratumor heterogeneity, cancer evolution, and therapeutic resistance. Subclonal diversification arises from genetic instability, selective pressures from the microenvironment and therapy, and epigenetic alterations that influence gene expression^[24]^. In the context of cancer therapy, treatment may selectively eliminate sensitive subclones while allowing resistant ones to persist or expand. This dynamic process underlies the acquisition of additional malignant features in response to therapy, either through the selection of pre-existing resistant clones or treatment-induced genetic changes^[25]^. Although intratumor heterogeneity has been widely implicated in resistance, the subclonal dynamics of ER-positive breast cancer during primary treatment remain poorly characterised. Gaining insight into how subclones are redistributed or reshaped during therapy at the personalised level—through the emergence or extinction of specific genomic alterations—could reveal resistance mechanisms and uncover new opportunities for therapeutic intervention.

In this study, we investigate mechanisms of endocrine response and resistance in locally advanced ER+ breast cancers, by characterizing the evolutionary dynamics of tumour subclonal architecture during sequential neoadjuvant AI therapy. This study builds on our clinical NeoLetExe trial^[9]^, which was designed to investigate the molecular basis of resistance to AI treatment by randomizing patients to receive letrozole followed by exemestane or vice versa^[9]^. We use whole-exome sequencing data from three timepoints in the NeoLetExe trial, to reconstruct cancer cell fraction– based subclonal composition and we validate subclonal structure with single-cell DNA sequencing from matched tumour samples. By tracking the clone trajectories during endocrine therapy, the study identifies mechanisms of resistance to the two aromatase inhibitors, including immune system involvement, and reveals therapeutic targets relevant to treatment failure.

## Materials and methods

### Patient Cohort and Study Design of the Neoletexe Trial

Patients diagnosed with locally advanced, ER+ breast cancer and eligible for neoadjuvant endocrine therapy were enrolled in NeoLetExe, a randomized, open-label phase-II trial^[9]^. Patients were randomized to receive monotherapy with either letrozole or exemestane for 3 months, followed by an intra-patient cross-over to the alternative drug for an additional 3 months. After completing at least 6 months of presurgical/neoadjuvant treatment, patients proceeded to breast surgery (breast conserving therapy or mastectomy) according to standard national guidelines. None of the patients received adjuvant chemotherapy. The study was approved by the Regional Ethics Committee of South-East Norway (project number 2015/84). All patients provided written informed consent for participation and for all procedures and analyses performed in this study. The trajectories of somatic variants detected in the patients across the three timepoints were followed and investigated on their involvement or association to AI resistance (see below). Patient IDs presented in this manuscript were randomly-allocated and are unrelated to the original study identifiers.

### Sample acquisition and handling

FFPE biopsies were taken before treatment initiation (baseline or Timepoint 1 (T1)), after 3 months of the first treatment, either letrozole or exemestane (crossover or Timepoint 2 (T2)) and after 3 months of the second treatment, either exemestane or letrozole (end of treatment or Timepoint 3 (T3)). All tumor biopsies were characterised by an experienced breast cancer pathologist at the Akershus University Hospital.

### Whole Exome Sequencing and RNA-sequencing

Ten 5-µm sections were prepared from formalin-fixed paraffin-embedded (FFPE) tumour blocks using standard microtomy. The uppermost section was stained with hematoxylin and eosin (H&E) and reviewed by a certified pathologist to delineate malignant regions. Guided by the annotated H&E slide, macrodissection was performed to isolate tumour-rich areas from the remaining sections. DNA and RNA were extracted from these regions using the Qiagen QIAamp DNA FFPE Tissue Kit and RNeasy FFPE Kit, respectively, and quantified with a Qubit fluorometer.

DNA libraries were constructed using the KAPA Hyper Prep Kit and enriched for exonic regions with the xGen Exome Research Panel v1.0 (Integrated DNA Technologies). Tumour and matched normal (germline) DNA samples were sequenced on the Illumina HiSeq platform, aiming for a coverage of ×200 and ×100, respectively. In selected samples used for clonality we performed higher than 200X sequencing coverage. For RNA-seq, libraries were prepared using the KAPA Stranded RNA-Seq Kit with RiboErase and sequenced to a depth of approximately 200 million reads per sample. RNA reads were aligned to the RefSeq (build 73) transcriptome using Bowtie2 (v2.2.6), and gene expression was quantified with RSEM (v1.2.25).

### Single Nucleotide Variation Calling

Single Nucleotide Variation (SNV) was performed using *GATK* (v4.1.8.1)^[26]^ to call SNVs in a (multi-sample when available) paired tumour/normal setting. In more detail, we first built a panel of normals (in Mutect2, germline samples are referred as normal) by running *Mutect2* on the 24 normal samples and merged germline information by running *GenomicsDBImport* and *CreateSomaticPanelOfNormals* in GATK, following the GATK guidelines^[27]^. We then, for each of the 24 cases, ran *Mutect2* in GATK on all of the available samples, considering the panel of normal to filter likely artefacts. Finally, somatic alterations identified by *Mutect2* were filtered using *FilterMutectCalls* in GATK and we discarded both non-A/C/G/T (removing indels & non-standard calls like multiallelic alterations) and SNVs not having a “PASS” filter. Somatic variants were detected using *SomaticSniper*^[28]^, *VarScan*^[29]^, *GATK*^[26]^, and *Pindel*^[30]^.

### Somatic variant detection

Structural variants (SVs) were detected with *BreakDancer*^[31]^. High-level analyses were performed with our *PathScan*^[32]^ and *MuSiC*^[33]^ packages, as well as *GeneGO MetaCore*^[34]^, and *PARADIGM*^[35]^. Candidate somatic variants and SVs were experimentally validated by custom capture and deep re-sequencing. This step was used for confirming predicted events and provided deep coverage for the reconstruction of tumour clonal architecture. Variant pathogenicity was inferred from ClinVar, classifying variants as pathogenic, likely pathogenic, variants of uncertain significance, benign and likely benign.

### Copy-number alteration (CNA) profiles

We extracted allele counts using *alleleCounter* (v4.0.0)^[36]^ at 29,076,189 SNP positions from the 1000 Genomes Project (GRCh37 phase 3, biallelic SNPs) across all samples, tumour and germline. SNPs with at least one read count in every sample were retained (n=921,657), and further filtered to those included in the whole-exome sequencing (WES) target design (n=226,774). Because allele counts varied between tumour samples (FFPE-derived) and germline samples, we applied additional filtering based on coverage distribution. We calculated the mean read counts for each SNP across germline samples, ranked them, and excluded SNPs with consistently extreme coverage (top and bottom 5%) to remove unreliable sites (n=204,096). The same filtering was applied independently to tumour samples, resulting in 183,686 SNPs retained. Following the methodology implemented in the *Battenberg* R package (v2.2.9)^[37]^, we derived logR and BAF values from allele counts, applying minimum count thresholds of 30 for germline and 1 for tumour samples. LogR values were corrected for GC-content and replication timing biases. Copy number analysis was performed using *ASCAT* (v2.5.2)^[38]^. For cases with multiple tumour samples, ASCAT was run in multi-sample mode using the *ASmultiPCF* algorithm^[39]^ to enhance detection of shared CAN, while for single tumour samples, the standard ASPCF algorithm was applied. Penalty and gamma parameters were set to 70 and 1, respectively.

### CNA filtering and clustering

To validate copy-number profiles, samples at baseline for each patient were harmonized for tumour ploidy, unless a CNA profile showed evidence of a different ploidy compared to other profiles for a given patient. Amongst the initial 56 samples, two samples were adjusted for ploidy, five samples were adjusted for purity and seven samples were adjusted for both ploidy and purity. Genomic changes were clustered using MEDDIC2 (based on 289 genomic segments larger than 100kb).

### Subclonal composition by DPClust

*DPClust* (v2.2.8)^[37]^ was used to cluster SNVs based on their cancer cell fraction (CCF), correcting VAF (from SNVs) for CNAs and tumour purity. CCF represents the fraction of tumour cells (CCF) that carry each somatic variant identified. Initially considering all 56 samples, we reran *DPClust* with refitted profiles and discarded samples below 20% tumour purity (28 samples from 11 patients). As a result of this filtering step, only five patients remained with three timepoints, while the remaining six patients had only two timepoints available. Samples surpassing the tumour purity threshold were used to infer tumour subclonal composition, by leveraging somatic variant cluster information, following general principles for reconstructing subclonal composition in tumour samples^[40]^. Small clones with <1% of SNVs were excluded from further analyses. CCF trajectories were tracked across timepoints and were categorised as persistent or eliminating after treatment, suggestive of resistance and sensitivity respectively. For each subclone, the mean cancer cell fraction at each timepoint was calculated as the average CCF of its constituent somatic variants. A mean CCF of zero at a given timepoint was interpreted as clonal disappearance, potentially indicating sensitivity to the treatment administered at that stage. In contrast, a stable or increasing mean CCF between consecutive timepoints was considered evidence of subclone persistence, suggesting persistence and potentially resistance to the treatment administered. For patients with only two timepoints, clones were still interpreted as eliminating or persistent, irrespective of the missing timepoint.

### Validation of identified clones by Mission Bio Tapestri

28 tumour Flash Frozen/Fresh Frozen (FF) samples from 11 patients from before were analysed using the Mission Bio Tapestri single-cell DNA sequencing platform to validate subclones identified by DPClust. Nuclei were isolated from biopsies using either the Singulator 100 system (S2 Genomics) or a manual protocol recommended by Mission Bio. A custom breast cancer panel targeting 528 patient-specific variants (497 amplicons) was designed based on previously identified mutations from WES. Due to costs constraints, not all variants assigned to DPClust-identified clones were included in the targeted panel. Libraries were prepared according to the Tapestri protocol and subjected to paired-end sequencing (150 bp) on an Illumina NovaSeq 6000 at the Norwegian Sequencing Centre (Ullevål Hospital). Sequencing data were processed with the Tapestri Pipeline, which included read alignment, cell barcoding, and variant calling. Downstream analysis was conducted using the *Mosaic* software and R. *Mosaic* was used to cluster variants into subclones, and assess clonal dynamics across treatment timepoints (T1–T3). Selected timepoints were analysed per sample (**Supplementary Table 1**), due to running costs or due to absence of FF tissue. Filtering steps included the removal of germline variants (>90% of cells), low-quality or low-confidence calls, and variants with inconsistent allele frequencies. Clone validation was defined as the presence of at least one variant from a DPClust-identified clone in the corresponding patient and timepoint based on scDNA-seq data.

### Variant functional annotation

All 1,928 variants detected in patient clones, where explored on their function, to distinguish functionally-important variants from passenger variants. Loss of function variants were predicted using Variant Effect Predictor (*VEP*) tool of Ensembl^[41]^. The impact of missense variants, was predicted using data from *Sift* (deleterious prediction), *Polyphen* (delta score≥0.85), *Alpha missense* (pathogenic prediction), *Revel* (score≥6) and *MetaRNN* (score≥0.8), based on which variants were annotated as pathogenic, disrupting protein function or deleterious. Splicing variants were then predicted using *dbscSNV* (ada score≥0.9 and rf score≥0.9), *MaxEntScan* (score≥2 or score≤-2) and *SpliceAI* (score≥0.5), identifying variants disrupt splicing at acceptor or donor sites or result in a change in canonical splice site strength. The machine-learning (ML) method, *Sei*^[42]^, was used to predict the functional impact of queried variants, based on chromatin profile enrichment (absolute z-score>1). Sei predicts the functional and regulatory impact of genetic variants based on ∼21,000 chromatin profiles across >1,300 tissues and cell lines. This machine learning framework assigns a functionality score to each variant and categorizes it into 40 tissue-specific or global regulatory mechanisms, which are referred as ‘sequence classes’ by the developers. To identify variants that directly affect gene expression, the *ExpectoSc* ML pipeline was applied (absolute z-score>1). *ExpectoSc* is trained on single-cell RNA-seq data^[43]^, in this case, based on a breast model^[44]^, leveraging cell-type–specific regulatory architecture.

### Gene annotation

Somatic variants detected in tumours, were annotated to genes by converging annotations from relevant sources. Firstly, coding, intronic and splice variants were annotated to the closest gene using the *VEP* tool^[41]^. Non-coding variants found within enhancers were further annotated using the activity-by-contact (*ABC*) model^[45]^, by taking the gene interacting with the enhancer based on Hi-C data from 12 breast-associated cell lines. Non-coding variants were also annotated through *ExpectoSc*^[43]^, a machine-learning pipeline trained on single-cell RNA-seq data from a breast model, that predicts expression disruptions from sequence at the cell-type specific context.

### Pathway Enrichment Analysis

To determine the enrichment of identified genes within pathways, also in association to resistance to AI inhibition, genes identified from before, were used as input to the functional module detection feature^[46]^ of the Human Base^[47]^, which performs a tissue-specific pathway enrichment analysis. The module detection feature, also separates pathways of similar function and involvement based on gene interactions in defined clusters.

### Association and Correlation analyses

A linear mixed-effects model was used to evaluate the association between variant clonality, measured by cancer cell fraction (CCF), and treatment response across timepoints. Fixed effects included CCF and timepoint, with a random intercept for each variant (VariantID) to account for repeated measures. The model was specified as: response ∼ CCF + timepoint + (1 | VariantID) and fitted using the lmer() function from the *lme4* R package. To examine whether the effect of CCF on treatment response varied across timepoints, a second model including an interaction term between CCF and timepoint was also fitted: response ∼ CCF * timepoint + (1 | VariantID). Treatment response was defined as the aberrant cell fraction estimated by ASCAT for each sample and timepoint.

## Results

### Somatic pathogenic variants in locally advanced ER positive breast cancers

Tumour biopsies were collected from 24 patients enrolled in the NeoLetExe trial from three timepoints: before treatment initiation, i.e. baseline (T1), after three months of first line AI treatment, either letrozole or exemestane (T2), and after three months of second line AI treatment, exemestane or letrozole, respectively (T3). All patients were postmenopausal. A subset of 58% of patients had grade II, 20.7% had grade I and only one presented with grade III. All tumours were ER-positive and HER2-negative, with low progesterone receptor expression (<10% of cancer cells) observed in four patients. Pathological assessment of residual disease at T3 relative to baseline (T1) revealed partial responses (>50% tumour reduction) in 22 patients, and no response (no tumour shrinkage / stable disease) in two. Other clinicopathological features are summarised in **Supplementary Figure 1**.

WES was performed on 56 tumour samples, resulting in the identification of 19,018 somatic variants (13,403 unique genomic locations). A summary of somatic variant frequency per patient is provided in **Supplementary Table 1**. On average, each biopsy harboured 340 somatic variants. One patient (L73) was a clear outlier, exhibiting a markedly elevated somatic variant burden. This was functionally supported by the presence of a pathogenic somatic missense *BRCA1* variant, consistent with a mutator phenotype^[48]^. Variant classifications were retrieved by ClinVar^[49]^, as follows bening, likely benign, pathogenic, likely pathogenic or uncertain (**Supplementary Table 2**). The majority of variants were classified as benign (82.7%, including likely benign). There were 519 pathogenic and likely pathogenic, associated with 227 genes. A histogram depicting the number of pathogenic or likely pathogenic variants found for the nine most frequently mutated genes are showin in **Figure 1A**. These were previously found mutated in luminal breast cancers treated with AI^[7,50]^. The most frequently mutated gene was *ZNF117* followed by *PIK3CA* and *PABPC1*. Results of Wilcoxon rank-sum tests on the association of genes carrying somatic pathogenic or likely variants to clinical parameters at each timepoints are showing in **Figure 1B**, including *PIK3CA* and Ki67 (lower in mutated tumours at timepoint T1), *CDH1* and ER status (lower in mutated tumours at timepoint T3), *MAP3K1* and PR status (lower in mutated tumours at timepoint T3) and *TBX3* with Ki67 (higher in mutated at T3) and magnetic resonance imaging (MRI, smaller size in mutated tumours) at T3. All data are shown in **Supplementary Figure 1**, where genes are represented as rows of a clustered variant matrix for the individual biopsies (columns) along with data on sequential order of AI treatment, tumour size measured by magnetic resonance imaging (MRI), and ER, PR and Ki67 staining from histopathological evaluation.

**Figure 1.**
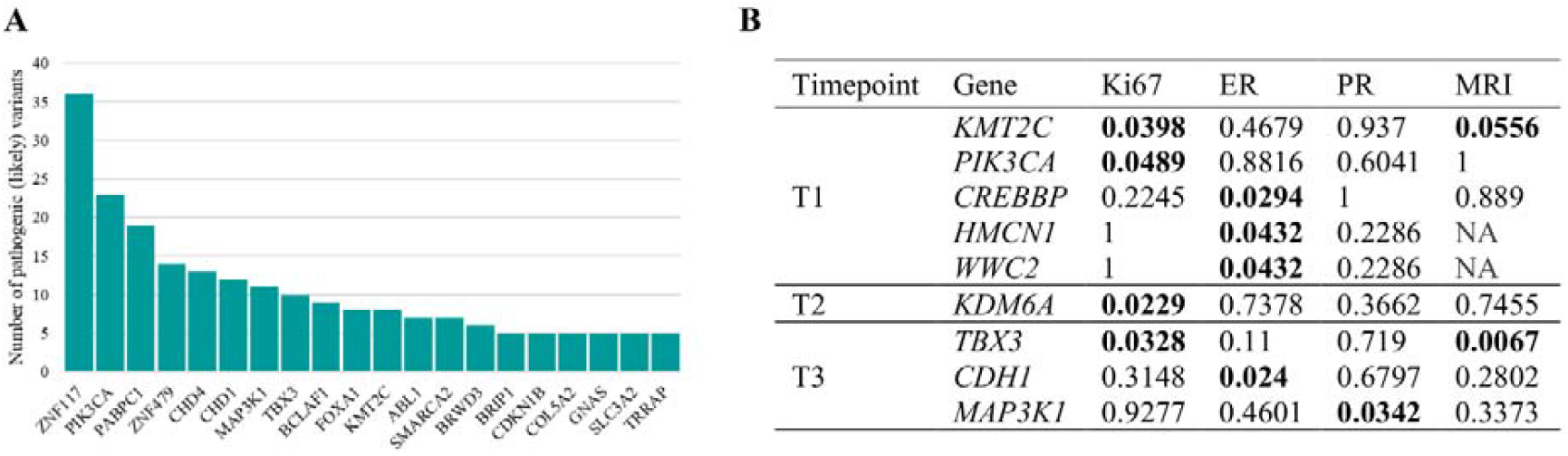
Pathogenic variants in locally advanced ER+ breast cancers. (**A**) Histogram showing the number of pathogenic or likely pathogenic somatic variants in genes with five or more such variants across the 56 exome samples. clinical parameters at each timepoints. Ki67, ER and PR were measured as percentages and MRI measurements in centimetres. Shown are genes with at least one significant result (highlighted in bold).

### Subclonal Dynamics Inferred from WES During aromatase inhibition treatment

Somatic variants and copy number alterations (CNA) were analysed in the 24 patients of the study. To discover shared CNAs across timepoints, CNA profiles from early tumour samples (T1 or T2) were compared to late samples (T3.1 or T3.2; **Supplementary Figure 2**). Recurrent alterations included gains in 1q (7/11 samples; 64%) and focal losses of chr19:9,045,897-9,237,147 (>80% of samples) encompassing genes such as *MUC16, OR1M1, OR7G1, OR7G2* and *OR7G3*. When analysing tumour purity, 11 patients were selected with ≥20% across all samples (**Figure 2A**). As a result of treatment, tumour purity was significantly lower in late samples, with median purities of 45.5%, 37.5% and 31.5% at T1, T2 and T3 respectively. The overall median purity across all samples was 39%. Analysis of subclonal architecture using DPClust revealed a strong positive correlation between somatic variant burden and subclonal diversity across the 11 patients (ρ = 0.79, P = 3.48×10□^3^; **Figure 2B**), indicating that higher mutation burdens facilitate more detailed subclonal resolution. Tumors with higher numbers of single nucleotide variants (SNVs) demonstrated increased subclonal complexity, suggesting that greater mutational load drives intratumor heterogeneity and clonal evolution in this patient cohort. A summary of measures obtained for these patients is provided in **Figure 2C**. Briefly, tumour ploidy is evident in specific samples (K12, M78, N29, E67), consistent with WGD events. As previously reported, patient L73 has the highest number of SNVs detected, representing an outline of the dataset. Based on the % inter-sample agreement (ISA), patients N29 and E67 show high subclonal heterogeneity between samples compared compared to the remaining samples that exhibit a consistent subclonal structure across timepoints (∼90-100 %ISA). Genomic instability (GI) also varies between samples and across timepoints. Higher GI score is observed in E67, followed by T1 in N29 and all timepoint samples of M78. Loss of heterozygocity (LOH) measurements across patients, are consisted with the accumulation of chromosomal alterations in all timepoints of E67, T1 of patient N29 and T1 and T3 of J47.

**Figure 2.**
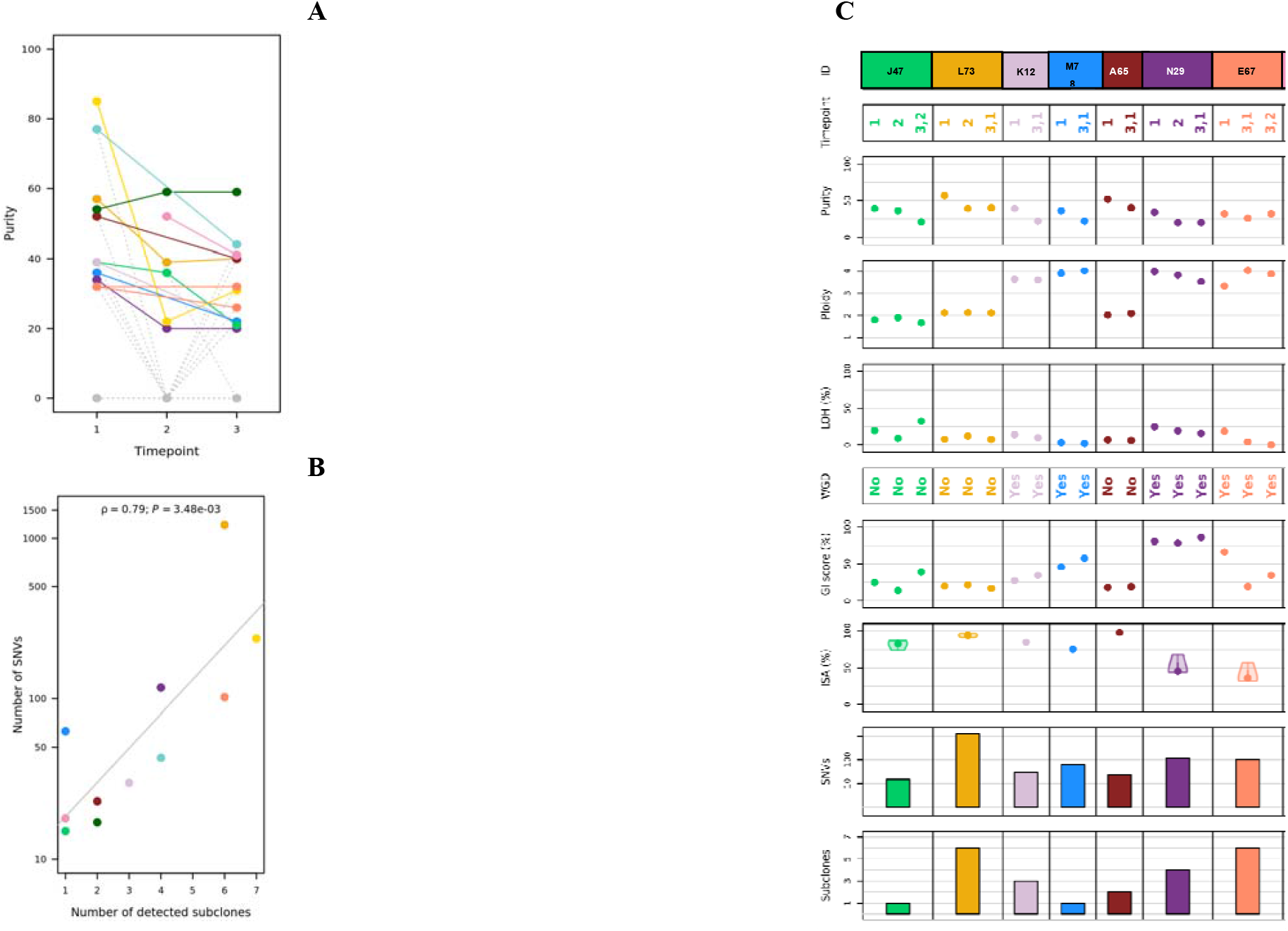
Tumour purity, relationshop between number of SNVs and subclones and sample characteristics. (**A**) Comparison of tumour purity at different timepoints (timepoint 3 combines 3.1 and 3.2 when available). Samples with less than 20% purity were set to 0 and dashed grey lines link such missing samples to the others, in respect to timepoints. In cases where T1 and T3 were available but T2 was discarded because of purity, a solid-coloured line directly links T1 to T3 (dashed grey lines still link 1 to 2 and 2 to 3). The dots are colour coded according to legend on top of panel C. (**B**) Correlation between number of detected subclones and number of SNVs. (**C**) Overview of purity, ploidy, % of LOH, WGD, GI score, % ISA, number of SNVs and subclones detected in each patient and timepoint (shown at top label). SNVs; Single-Nucleotide Variants, LOH; Loss of Heterozygocity, WGD; Whole-genome duplication, GI; Genomic instability, ISA; Inter-sample agreement

DPClust-derived subclonal evolutionary trajectories and clonal dynamics were tracked in response to treatment over time. Results for patients L73, N29, and V27 are shown in **Figure 2A-C**. For each patient and timepoint, cancer cell fractions (CCFs) are displayed for subclone-associated variants (blue heatmap panel), representing the proportion of tumour cells harbouring each subclone. Subclonal phylogenetic trees are also shown, along with a 100-cell visual representation depicting the hierarchical structure of clones. Briefly, in L73 (assigned in the letrozole-exemestane arm), a total of six subclones were detected by DPClust of variable evolutionary patterns (**Figure 2A**). Conversely, in N29 (**Figure 2B**), four subclones are detected, of which their representation based on clone and variant CCF remain constant over time, indicative of treatment resistant to both treatments. Specifically, two clones emerge in T2 as a response to letrozole, indicative of treatment resistance, with one of them remaining present also in T3, indicative of resistance to both treatments. Additionally, two clones persist throughout timepoints, potentially indicative of resistance, one clone is only present at baseline, indicative of sensitivity to both letrozole and exemestane, while one clone emerges in T3 as a response to exemestane. Similarly, the subclonal reconstruction of V27 identified five distinct clones. **Figure 2C** shows their temporal presence of seven clones: three clones present at baseline, regress at T2, indicative of sensitivity to letrozole, which reappear in T3 after exemestane, indicative of lack of cross-resistance. One clone persists throughout the timepoints, suggestive of resistance to both treatments, two clones appear in T2, indicative of letrozole resistance, and one clone appearing at T2, persists through T3, suggestive of resistance to both treatments. Any remaining patient samples are presented in **Supplementary Figures 3-10**.

**Figure 3D** provides an overview of the diverse clonal trajectories observed across timepoints and patients in the study. The mean CCF of each clone was used to determine evidence of clone persistence (CCF_mean_>0) or elimination (CCF_mean_=0) across timepoints. Clones eliminated are represented by grey circles, indicative of potential sensitivity to the treatment administered at each timepoint. Persistent clones (whether stable or expanding) are represented by colour-coded circles, and were considered as potentially resistant to the treatment administered. Missing timepoint samples discarded due to low purity are represented by white circles. Variable clone trajectories are observed across timepoints, including the re-emergence of clones following a specific treatment, persistent clones across treatment intervals, eliminating clones suggestive of sensitivity, and newly emerging clones likely induced by therapy. Based on the available timepoints, the treatment and sequence of treatment administered, and the CCF dynamics, subclones were classified into the following categories: (i) clones persistent after both treatments, (ii) clones persistent after letrozole or exemestane (for patients with two timepoints), (iii) clones persistent after exemestane and eliminated after letrozole, (iv) clones persistent after letrozole and eliminated after exemestane, (v) clones eliminated after both treatments, (vi) clones eliminated after letrozole or exemestane (for patients with two timepoints). **Figure 3E** shows the distribution of identified clones (n=60). Categories presenting with evidence of lack of cross-resistance (i.e., persistent to one treatment and disappearance after another) were further subdivided according to the order of drug sequence. Clones persistent after both treatments (irrespective of drug order) were the most common category, accounting for 33.3%, followed by clones that persist specifically after exemestane (26.7%), identified in patients with two available timepoints. All remaining categories ranged from 1 to 5 clones, in which 5% were eliminated after exemestane, 1.7% were eliminated after letrozole and 1.7% were eliminated after both treatments. Together clones under lack of cross resistance represent 25% of clones detected. **Figure 3F** shows the distribution of different clone categories in patients, revealing that persistent and eliminating clones frequently co-occur within the same patient, highlighting the intratumoural heterogeneity and differential clonal responses to treatment. Among the variants and clones identified by clonal reconstruction, some were present at baseline (T1) before treatment started, while others emerged as a response to treatment. Specifically, out of the 60 persistent clones with CCF measurements in T1, the majority of clones (63.3%) were already present at baseline, likely reflecting intrinsic resistance, whereas 23.3% emerged after treatment initiation, potentially indicating acquired resistance.

**Figure 3.**
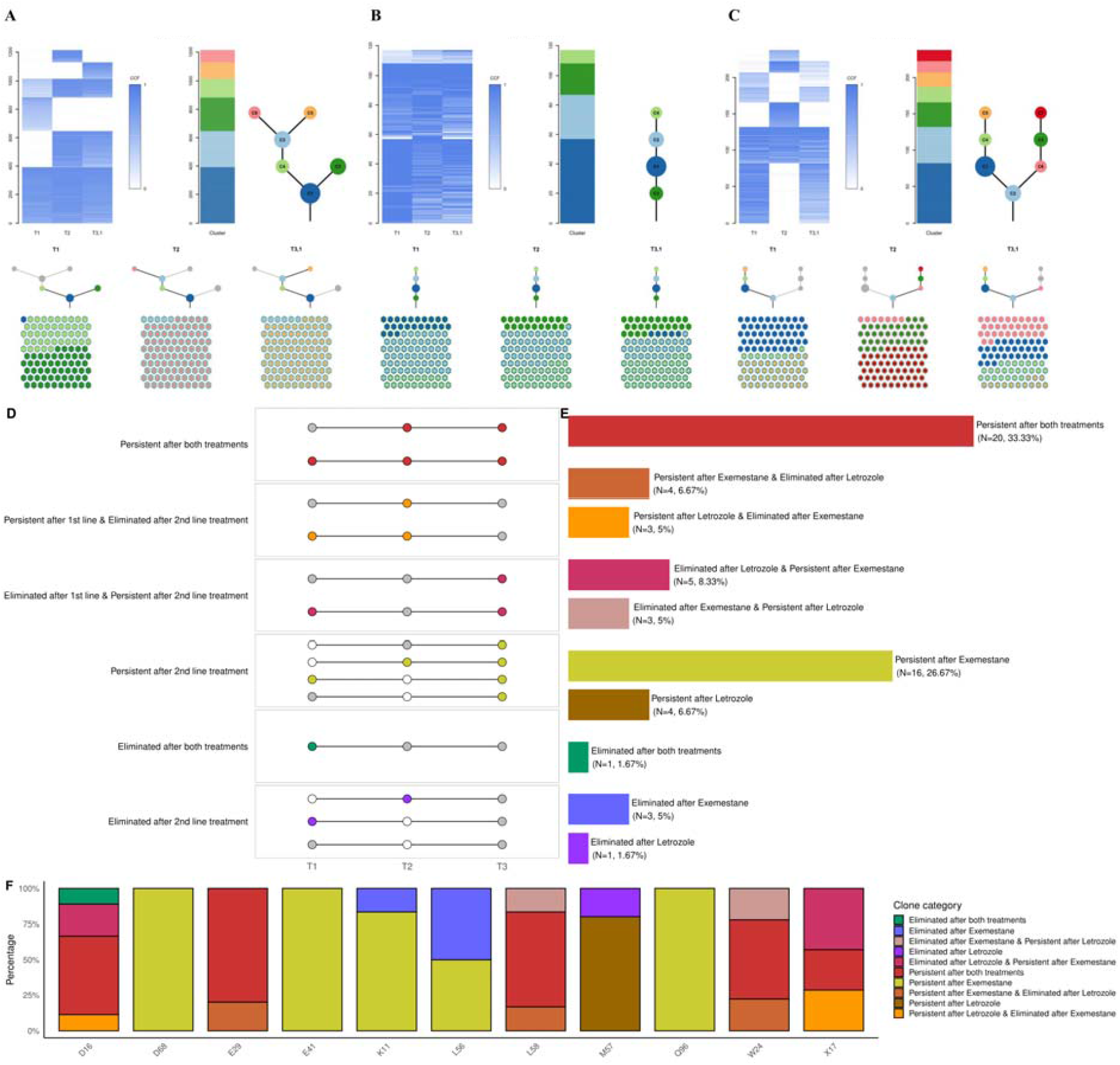
Example representation of DPClust output and summary of clonal drynamics identified in patients. DPClust outputs for different patients: (**A**) L73, (**B**) N29, (**C**) V27. The blue histogram panels represent variant CCF values in clones (y-axis) at each timepoint (x-axis). In each figure, a phylogenetic tree of the subclones (in colour) detected in each tumour are provided, depicting the evolutionary relationships between them. A representation of 100 cancer cells with nested colours are also provided in the bottom panels, indicating the presence of evolutionary sweeps during treatment throughout the three timepoints of the trial. (**D**) Summary of clonal trajectories across the three timepoints, T1, T2 and T3. Clone status is colour-coded: white indicates clone absence due to unavailable sample (low tumour purity), grey denotes true clone absence (CCF_mean=_0), and coloured circles represent clone presence (CCF_mean_>0) grouped according to persistence or disappearance after either or both treatments. (**E**) Subclassification of clones into distinct categories based on treatment type and sequence. The number and % of clones identified per category is provided in the brackets. (**F**) Distribution of clone categories per patient. Clone categories are consistently color-coded across panels.

CCF clone trajectories were compared to ASCAT-derived aberrant cell fractions, considered a reliable response variable. A linear mixed effects model adjusting for variant-level random effects revealed a significant negative association between CCF (raw CCF) and ASCAT response (β = – 1.45, p = 9.5×10 □ □), suggesting that higher clonal cell fractions may be linked to reduced treatment response. Timepoint-specific effects were modest, with only T2 showing a borderline increase in response (p = 0.023), while T3 was not significantly different from baseline. Additionally, a linear mixed effects model with interaction terms revealed that higher CCF (raw values) was significantly associated with reduced response at baseline (T1, β = –2.40, p < 0.0001). This negative association was significantly attenuated at T2 (CCF × T2 interaction, p = 0.005), while the effect at T3 was weaker and not statistically significant (p = 0.20). These findings suggest that the relationship between subclonal burden and treatment response may be modulated by treatment phase.

### Single-cell DNA sequencing validation of the subclonal architecture of the tumours

Somatic variants identified through WES and assigned to clones via DPClust were used to design a targeted scDNA-seq panel on the Mission Bio Tapestri Platform to evaluate tumour subclones. **Supplementary Table 3** details the number of variants, clones and timepoints assessed per patient. **Figure 4A** summarizes the panel design, including the number of variants, amplicons and clone specific variants selected for Mission Bio validation. **Figure 4B** presents validation results stratified by patient and clone. While variant-level validation averaged 67% per patient, clone-level validation achieved 81% across the cohort. Three patients demonstrated complete clone validation (100%), with the remaining patients achieving validation rates between 50–89%.

**Figure 4.**
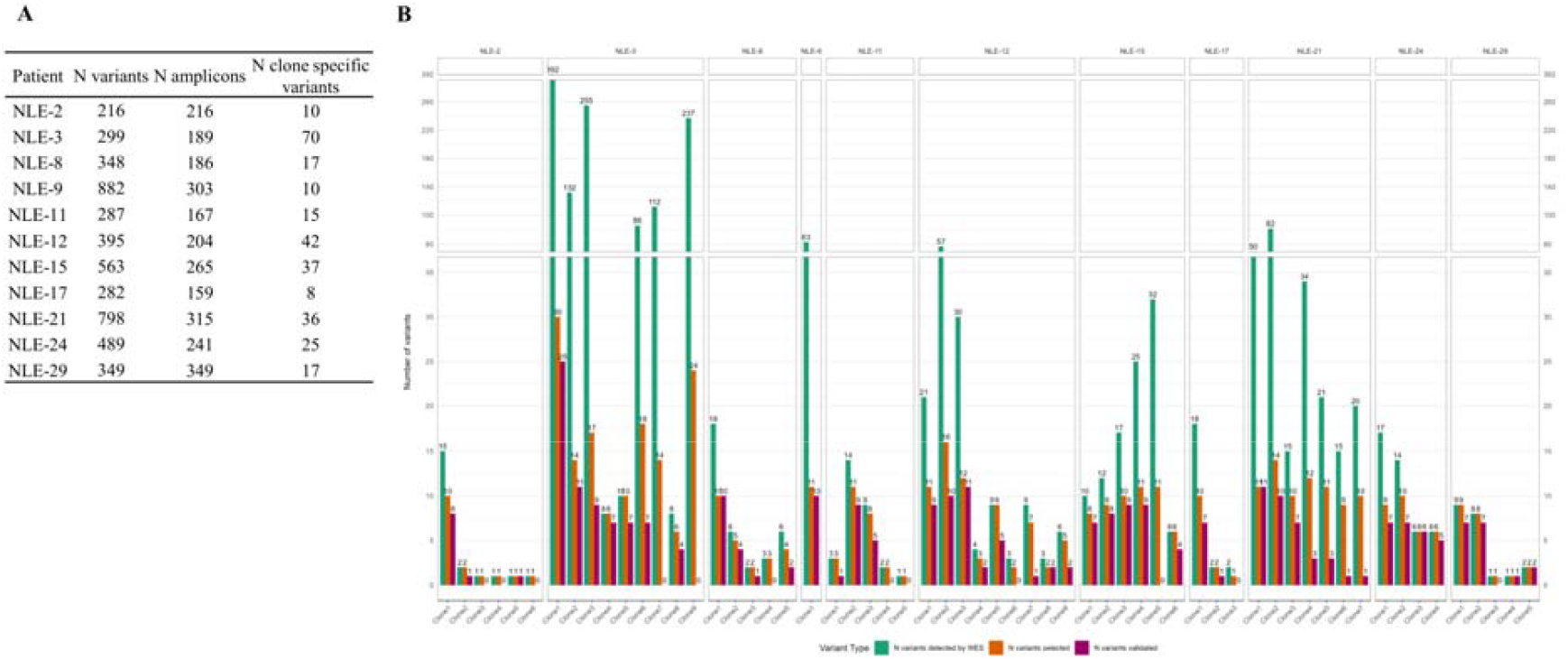
Validation of clones and variants detected by DPClust using Mission Bio. (**A**) Summary of the number of variants, amplicons and number of clone specific variants subjected to scDNA sequencing in the same samples. (**B**) shows a summary of the Mission Bio results per patient (facets) and per clone (x-axis), as well as the number of variants (y-axis) detected by WES in the same samples (orange bars), those selected for scDNA sequencing by Mission Bio (green bars), and those successfully validated (pink bars).

### Sublocal Variant Characterisation and Key Genes Modulated

Somatic variants assorted in clones included non-synonymous (67%), synonymous (26%) and stop-gain variants (6%), with other consequences representing <1% (**Figure 5A**). Variants were further categorised according to functional consequence into single-cell eQTLs (8%), regulatory variants (3%), impactful missense variants (39%) and splice variants (1%) (**Figure 5B)**. No functional consequence was identified for 43% of variants. Subclonal variants exhibited pronounced intra-tumour heterogeneity, with only three shared across patients (**Supplementary Figure 11**), annotated to the genes *SF3B1, PIK3CA*, and *RBMX*. Consequently, detecting variants as universal markers of resistance was not feasible. Suclonal variants were also characterised on their ClinVar class, in which the majority of them had not been reported before (**Figure 5C**).

**Figure 5.**
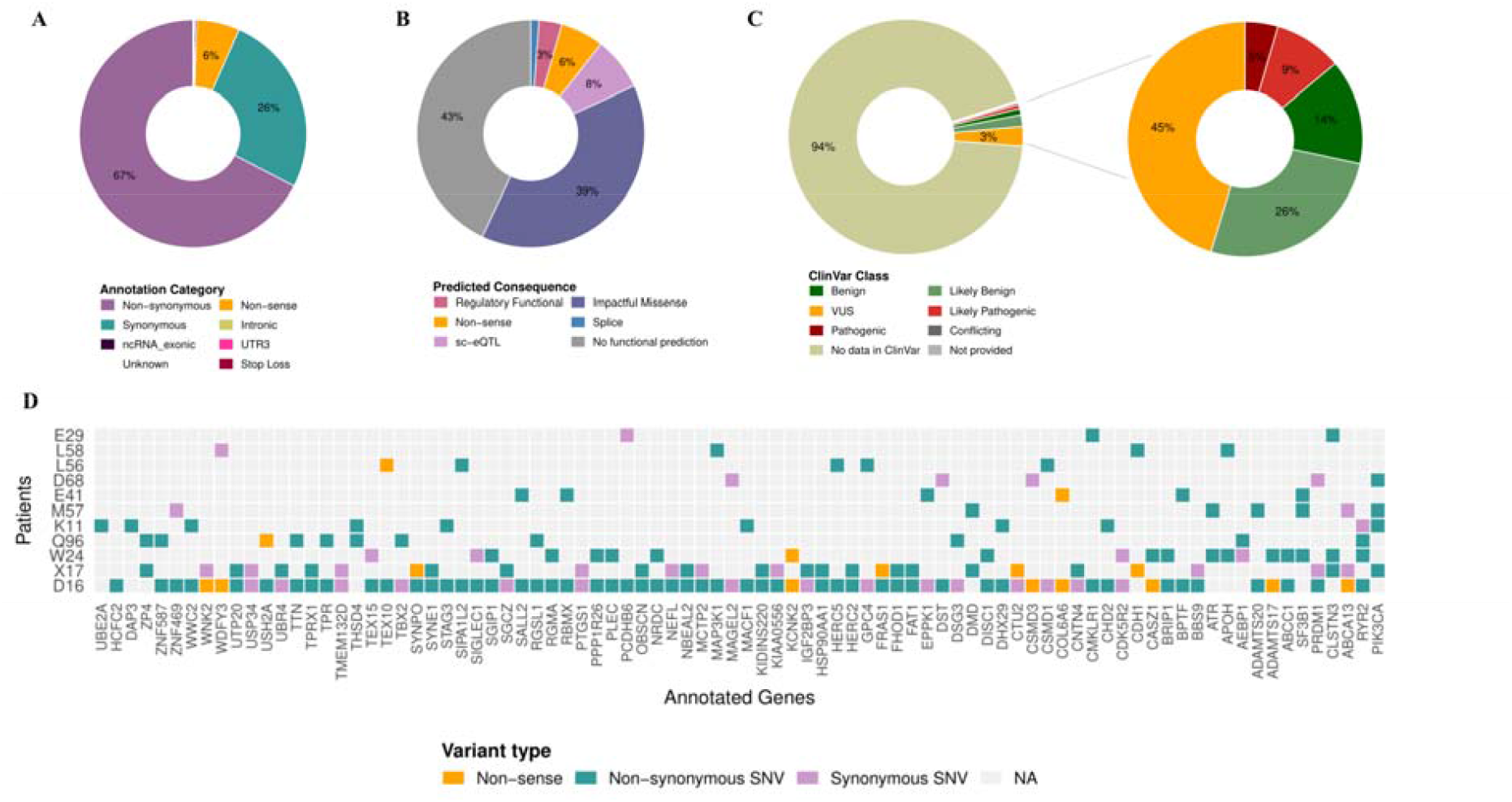
Variant characterisation and common genes annotated to subclonal variants. (**A**) Donut plot showing the distribution of variant consequence according to the closest annotated gene. (**B**) Donut plot showing the distribution of variant functional impact, including effects on gene expression, regulatory potential, impactful missense consequence and splice effects. (**C**) Donut plot showing the distribution of ClinVar class, separated into two panels; first panel: ClinVar class annotations, second panel: ClinVar class of benign, likely benign, VUS, pathogenic and likely pathogenic. (**D**) Histogram of common genes (x-axis) annotated to subclonal variants identified in each patient (y-axis). Histogram tiles are colour coded according to variant consequence. sc-eQTLs; single cell expression quantitative trait loci.

To investigate the biological relevance of clonal trajectories during sequential AI treatment, variants were annotated to target genes using a custom prediction pipeline, integrating evidence from difference sources and methods. Across the total 1,701 unique genes annotated to subclonal variants, 88 were found common between at least two patients, dropping to 15 after exclusion of outlier patient D16 (**Supplementary Figure 12A**). **Supplementary Figure 12B** shows gene overlaps across clone categories. Notably, most genes within each clonal category were unique, reflecting further intra-clonal heterogeneity, with only 91 targets annotated to more than one clone category. **Figure 5D** summarizes variant consequences within common annotated genes. While no consistent gene has emerged in potentially resistant or sensitive clones, suggesting patient-specific mechanisms, notable overlaps included *PIK3CA, CDH1, APOH, DAP3*, as well as *DMD, ABCA13, CLSTN3, ZP4, THSD4, RYR2, AEBP1* and *PRDM1*.

Among the genes disrupted by subclonal somatic variants were genes annotated to pathogenic or likely pathogenic variants in bulk WES (**Figure 1A**), including *PIK3CA, CDH1, GATA3, MAP3K1, SF3B1, TBX3, WWC2, CHD4, KMT2C, ABL1, SMARCA2, BRWD3, BRIP1, COL5A2, GNAS*, and *TP53*. Other genes annotated to pathogenic or likely pathogenic variants included *FGFR2, LRP5, ATM, AKT1, SF3B1, SNORA4, COL4A4, RTEL1, PHIP*, and *BRAF*. Two variants of uncertain significance (VUS) were annotated to *BRIP1*. All remaining variants with a reported ClinVar class were not associated with any annotated gene. Among the genes annotated to subclonal variants included 57 known cancer drivers, three which were annotated to non-sense variants, 11 to synonymous and 43 to non-synonymous changes. **Supplementary Figure 13** shows the CCF trajectories of 56 subclonal variants present at baseline, indicating intrinsic resistance, as well as the trajectories of 17 subclonal variants emerging as a response to either letrozole or exemestane, indicative of acquired resistance. Variants were stratified by the annotated driver genes. ExpectoSc identified 157 predictions subclonal variants with cell-type specific variant-gene effects in stromal, endothelial, breast and immune cell types, suggesting functional compartmentalisation of regulatory impact within the tumour microenvironment (**Supplementary Figure 14, Supplementary Figure 15**). Key genes found upregulated or downregulated by clone variants are shown in **Figure 6**. *SMARCA1* was predicted to be strongly downregulated (Z-score = –176.0) and *RBMX* markedly upregulated (Z-score = 225.9) in luminal epithelial cells. Clones persisting after letrozole carried baseline variants in drivers such as *ATR, PIK3CA*, and *SF3B1*, while persistence after exemestane involved emerging variants in *SF3B1* and *RBMX*. Intrinsic drivers including *ABL1, AKT1, KMT2D, PTCH2, JAK2*, and *THRAP3* were disrupted through missense, nonsense, or regulatory changes. Genes such as *CBLB, KDM5C, ARID1A, BRAF, FAT1*, and PIK3CA displayed selective persistence under one AI but loss under the other, reflecting treatment-specific selective pressures. Two additional drivers, *SPEN* and *CHD4*, emerged following exemestane at T2 but disappeared after subsequent letrozole exposure, whereas a *FGFR2* missense variant persisted under letrozole but was eradicated after exemestane. Finally, several canonical drivers—including *FAT1, JAK1, ARID1B*, and *ATM*—were linked to clones eliminated after treatment, consistent with a role in sensitivity rather than resistance. The overall ExpectoSc zscore distributions of identified predictions are shown in **Supplementary Figure 16**, in which clone variants persisteny to both treatments or after exemestane exhibited significantly higher effects compared to clone variants eliminated after both treatments or those lacking cross-resistance.

**Figure 6.**
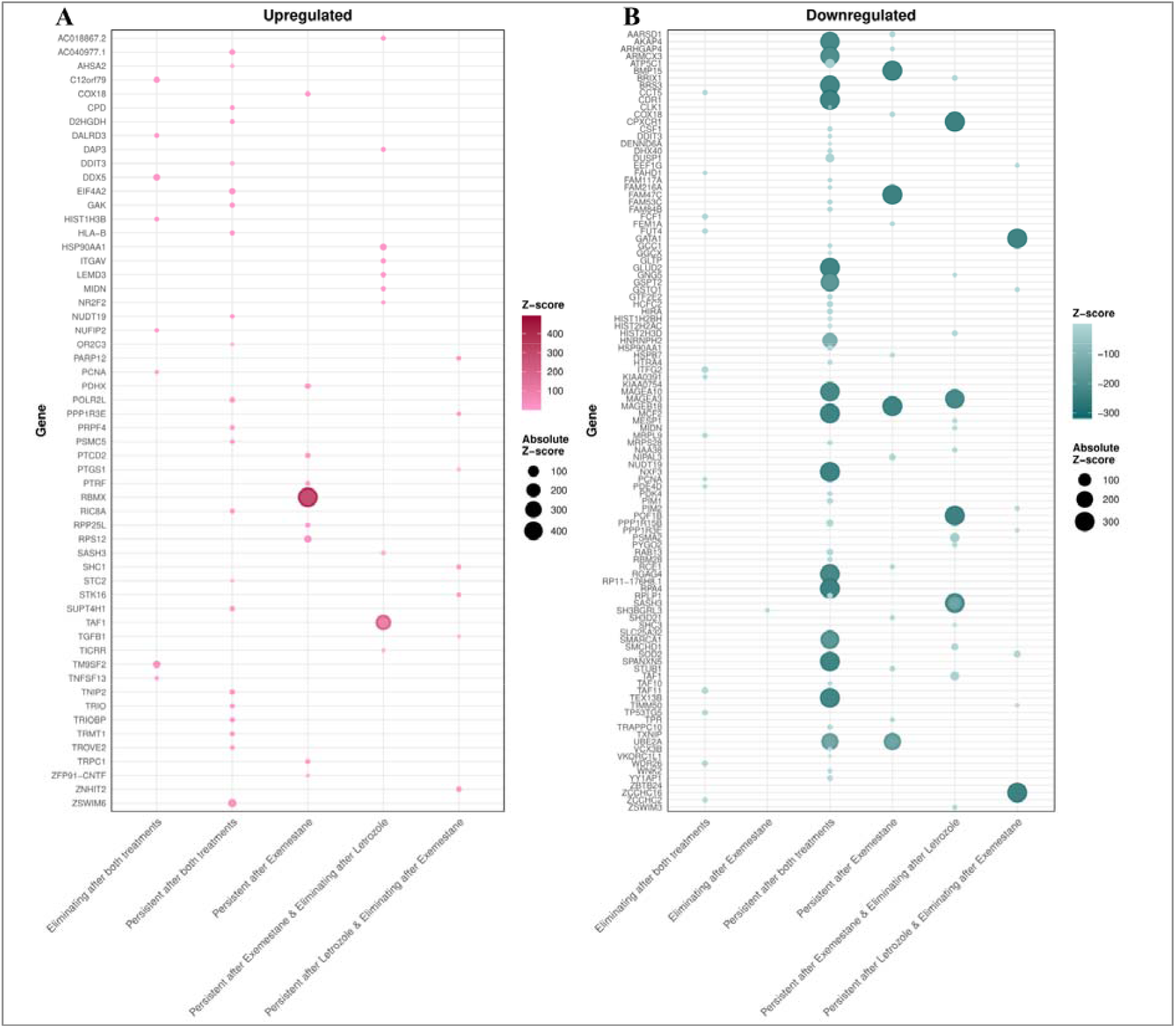
ExpectoSc-predicted gene expression changes across clone groups. Bubble plot size and colour represents the z-score expression change for each gene (y-axis) for each clone group (x-axis). Genes predicted are separated into upregulated and downregulated separately.

### Mechanisms and Pathways of resistance and sensitivity to Aromatase inhibition

Annotated genes were used in pathway enrichment analysis, stratified by clone categories, to identify mechanisms and pathways involved in resistance and sensitivity to AI treatment. The Human Base tool was used for this purpose which integrates pathways from Gene Ontology, MSigDB Hallmarks, and MSigDB C2. Pathway co-occurrene between clone categories are presented in **Supplementary Figure 17**. Cell-matrix adhesion, mitotic spindle and downregulation of the KRAS signalling were commonly observed pathways between clone categories. Among the pathways shared between two clone categories were: TP53 and MYC pathways, IL2 STAT5 signalling, response to tumor necrosis factor, hypoxia, epithelial mesenchymal transition, histone H4-K16 acetylation and H3-K4 methylation, production of miRNAs involved in gene silencing and fatty acid metabolism. A curated selection of pathways at *P*-adjusted < 0.01 stratified by clone category are shown in **Figure 7**.

**Figure 7.**
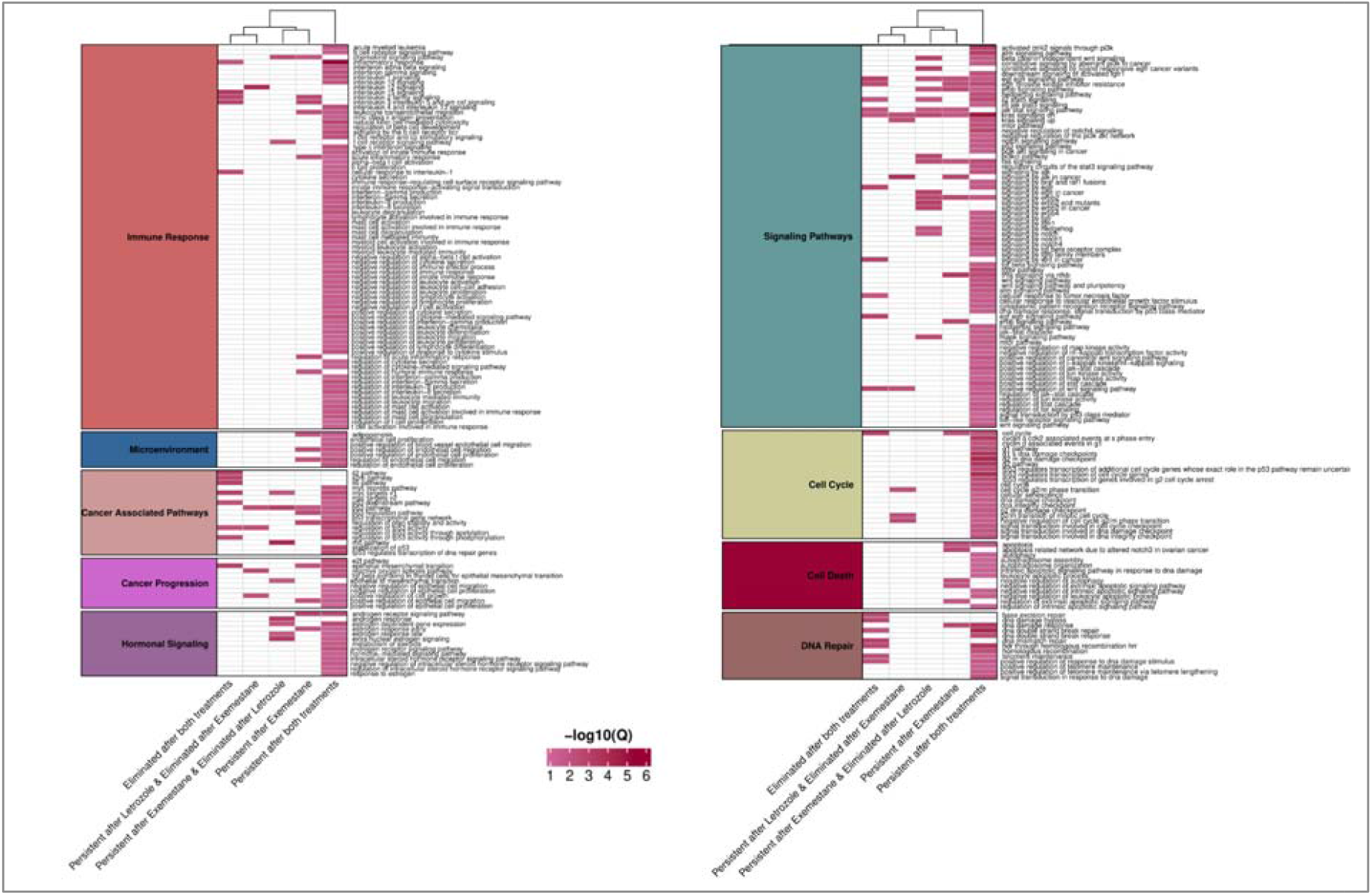
Pathway enrichment analysis stratified by clone categories. Heatmap displaying pathway enrichment results obtained using Human Base for genes within each clone category. Colour intensity represents the strength of enrichment (-log10 Q-value), with darker colours indicating higher significance. Only pathways with Q-values < 0.01 are displayed. Pathways are grouped by related biological processes, as indicated by the coloured annotation bars.

#### Pan-Resistant Clones (Both Letrozole and Exemestane)

Clones persistent to both aromatase inhibitors demonstrated the most complex profile, with extensive pathway enrichment spanning six major categories. Epigenetic reprogramming was prominent, including chromatin remodeling, histone modifications (H3-K9, H4-K16 acetylation), and pre-miRNA processing. DNA damage response pathways were consistently enriched, encompassing G2/M checkpoint regulation (*BRCA1, CDC25C*), TP53-mediated signaling, hallmark gene sets like E2F targets and double-strand break repair mechanisms. Notably, immune activation pathways showed extensive enrichment, including interferon responses (*EPSTI1, IFI44*), complement activation (*CFH, C3*), mast cell degranulation and activation (*ADGRE2, CD300A, KIT*), JAK-STAT signaling, TNFα/NFκB pathways (*DUSP1, CSF1, ICAM1*) and cytokine secretion. Both innate and adaptive immune pathways were represented, including toll-like receptor signalling, IL6-JAK-STAT3 signalling (*CSF1, PIM1*), TNFα signalling via NFκB (*DUSP1, ABCA1, CSF1, ICAM1*), and myeloid leukocyte activation, migration and degranulation (e.g. *ADGRE2, KIT*). Metabolic reprogramming was also evident through fatty acid oxidation, glycolysis, and mitochondrial reorganization pathways. Cellular plasticity mechanisms included epithelial-mesenchymal transition (EMT; *COL5A2, BDNF*), cell migration, and angiogenesis pathways. Most relevant to AI resistance, steroid hormone signaling was involved through estrogen response pathways (*RBBP5, EP300*), androgen receptor signaling (*MED13, EP300*), negative regulation of steroid hormone receptor signalling and pro-growth signals including KRAS, WNT (*USP34, LRP6*), mTOR signaling (ARAF, WDR24) and hedgehog signalling (*CELSR1, GLI1*). Other important pathways highlighted include angiogenesis (*AGTR1, ANXA1*), apoptosis (*BRCA1, VDAC2, DAP3*), adipogenesis (*LAMA4, ABCA1, C3*) and autophagy regulation.

#### Drug-Specific Resistance Patterns

Exemestane-only resistance was characterized by distinct telomere maintenance pathways (*EXOSC10, TENT4B*), RNA processing mechanisms, and acute inflammatory responses (*CR1, NLRP3*). Apoptosis regulation (*BIRC6, PHIP, DAP3, TXNIP*) and oxidative stress responses were prominent, alongside cellular adhesion and migration pathways. Other statistically significant pathways included the regulation of acute inflammatory response (*CR1, NLRP3*), humoral immune response (*PGC, CR1*), response to tumor necrosis factor (*MAP4K3, JAK2*), and interferon gamma response (*TXNIP*). Clones also exhibited upregulation of ubiquitin-related and oxidative stress-related processes (e.g., cellular response to reactive oxygen species), as well as pathways related to cellular adhesion (*MACF1, ITGA4*) and migration of endothelial and epithelial cells (*ABL1, AKT1*). Metabolic and signalling signatures such as adipogenesis (*SCARB1, CAVIN1*), early estrogen response, P53 pathway, complement, hallmark hypoxia, and TNFA signalling via NFkB were also found significantly enriched.

Cross-resistance patterns revealed distinct mechanistic profiles. Exemestane-resistant/letrozole-sensitive clones demonstrated epigenetic control through histone modifications (H3-K4 methylation/trimethylation) and miRNA-mediated silencing (*PUM2, AGO2*), retaining complement pathways (*PIK3CA, CBLB*) and hormone responses including androgen response (ABCC4, ITGAV, IDI1), late estrogen response (PRKAR2B, TIAM1). Other pathays strongly involved included MYC targets (*PSMA2, UBA2*), P53 Pathway (*PMM1, RETSAT, PLXNB2*), downregulation of KRAS signalling (*SYNPO, TENM2, RYR1*), IL2-STAT5 signalling. Critically, letrozole-resistant/exemestane-sensitive clones showed metabolic focus (fatty acid metabolism, IL2-STAT5 signaling) and cell cycle control but lacked immune pathway activation, suggesting exemestane effectiveness correlates with immune pathway suppression. Late estrogen response (*PRKAR2B, TIAM1*) and androgen response (ABCC4, ITGAV) were also prominent.

#### Pan-Treatment-Sensitive Clones

Pan-sensitive clones maintained coordinated cell cycle control (E2F targets, G2/M checkpoint), positive DNA damage responses (*PCNA, DDX5*), and crucially, preserved immune signaling (IL2-STAT5, TNF/interleukin responses). The retention of stress-response pathways (Wnt, hypoxia, KRAS signalling (downregulation), UV response) in sensitive clones contrasts sharply with resistance mechanisms, indicating that pathway silencing rather than activation drives treatment failure. EMT-related (epithelial-mesenchymal transition) and MYC-regulated (MYC targets) gene sets were also among the enriched terms. No significant pathways were found enriched to subclones eliminated by either letrozole or exemestane alone.

#### Clinical Implications

Analysis of FDA-approved breast cancer targets revealed actionable alterations for alternative or combinatorial therapeutic options. These included, the targets *PIK3CA* (4 patients), *AKT1* (1 patient) were disrupted by likely pathogenic or pathogenic subclonal variants. As presented before, the somatic variant affecting *AKT1* persisted after exemestane, indicating potential resistance. RNA-expression data on *AKT1* also revealed consistently high expression (>90% TPM percentile) across timepoints (TPM_T1_=37.34, TPM_T2_=23.24). Similarly, the two disruptive subclonal variants in *PIK3CA*, affecting four patients, persisted after exemestane and letrozole, while it also regressed as a response to letrozole and re-emerged again after exemestane. *PIK3CA* expression levels, although below the 90^th^ percentile cutoff, showed minimal changes between timepoints (TPM_T1_=5.56, TPM_T3_=6.95 for NLE-21, TPM_T1_=9.18, TPM_T3_=7.30 for NLE-15 (correlating with CCF changes), and TPM_T1_=11.1, TPM_T3_=5.74 for NLE-11 (correlating with CCF changes)). RNA expression analysis identified 48 additional therapeutic targets spanning PI3K/AKT/mTOR, CDK4/6, DNA repair (BRCA1/2, ATM), and immune checkpoint pathways, providing a foundation for combination therapeutic strategies targeting resistant subclones. *ATM, AKT1, RPS6KB1* and *BRAF* were among the genes annotated to by subclonal variants, while ATM expression levels in tumours correlated with CCF trajectories.

## Discussion

In post-menopausal ER+ women with breast cancer, approximately 10-20% relapse during or after adjuvant endocrine therapy^[51]^, while virtually all patients with ER+/HER2-metastatic breast cancers ultimately develop resistance^[52]^. Key endocrine agents include aromatase inhibitors, yet resistance—whether intrinsic, driven by tumour characteristics, or acquired through adaptation— remains a major clinical challenge^[53]^. Although diverse variants converge on similar deregulated pathways implicated in AI resistance, pathway-targeted therapies are lacking. The NeoLetExe trial was designed to explore the biological mechanisms of switching steroidal (exemestane) to non-steroidal (letrozole) AI treatments, and to investigate the molecular basis of cross-resistance.

Building on our prior work^[9–23]^, we performed whole-exome sequencing (WES) of tumours from 29 patients treated sequentially with exemestane and letrozole (in either order), based on which subclonal architectures were later reconstructed. WES identified 519 ClinVar-classified pathogenic or likely pathogenic variants. The most frequently affected genes were *ZNF117, PIK3CA*, and *PABPC1* (**Figure 1**), with *PIK3CA*, a central component of the PI3K/AKT pathway, highlighted for its established role in tumourigenesis and AI resistance^[50]^. Additional pathogenic variants were observed in genes previously linked to luminal breast cancers treated with AIs^[7]^, including *CBFB, CDH1, CDKN1B, GATA3, MAP3K1, SF3B1, TBX3*, and *TP53*, reinforcing their relevance to endocrine resistance. Exploratory clinicogenomic analyses revealed that *PIK3CA* variants associated with lower Ki67 at T1, suggesting a link between *PIK3CA*-driven tumor biology and lower proliferative indices at early treatment stages. *CDH1* pathogenic variants were also associated with lower ER expression at T3, and *MAP3K1* with lower PR status at T3, further supporting the potential role of these genes in the modulation of hormonal receptor expression during AI therapy. *TBX3* pathogenic variants were associated with higher Ki67 and smaller MRI-measured tumour size, suggesting an aggressive yet smaller-volume phenotype early on treatment. These finding pave the way for personalized therapeutic strategies in the treatment of endocrine-resistant breast cancer.

Tumours comprise multiple genetically distinct subclonal populations, potentially with unique responses to treatment. Although the influence of subclonal variation on treatment response and resistance is well documented^[25,54–56]^, its role in AI resistance remains incompletely defined.To dissect intra-tumour heterogeneity during sequential AI therapy, we applied DPClust to 11 patients with sufficient tumour purity to reconstruct subclonal populations. Subclonal pressures were tracked using evolutionary phylogenetic trees and CCF trajectories (**Figure 3**). We observed diverse patterns, with clones classified as persistent or eliminated (**Figure 4**). Specifically, 33.3% persisted after both treatments, 26.7% after exemestane, and 6.7% after letrozole, indicating more clones showing evidence of resistance following exemestane compared with letrozole. Also, 1.7% of clones were eliminated after both treatments, 1.7% after letrozole and 5% after exemestane. Clones that persist in response to one treatment but eliminated after the second line of treatment accounted for 25%, while 63.3% of persistent clones were already present at baseline, consistent with intrinsic resistance.

To validate subclones and variants detected by DPClust, we designed a targeted scDNA-seq panel on the Mission Bio Tapestri platform. Validation averaged 67% of clone-associated variants per patient and 81% overall, with 3 patients at 100% and the remainder 50–89% (**Figure 2**). These results support the accuracy and feasibility of WES-guided subclone inference for treatment-response studies.

CCF trajectories were assessed in relation to ASCAT-derived aberrant cell fractions across timepoints. Raw CCF showed a strong negative association with ASCAT response (β = –1.45, p = 9.5×10□□), indicating reduced response at higher clonal burden. Timepoint effects were modest: a borderline increase in response at T2 (p = 0.023) and no change at T3. At baseline (T1), higher CCF associated with lower response (β = 2.40, p < 0.0001), with attenuation at T2 (p = 0.005) and loss at T3 (p = 0.20). These findings suggest that the impact of subclonal burden is greatest at baseline and diminishes as treatment reshapes clonal architecture to timepoint 3.

Our analysis of subclonal variants showed variable annotation categories and characteristics (**Figure 5**) and highlighted the extensive genetic heterogeneity underlying AI resistance. Although variant types were diverse, only three genes (*SF3B1, PIK3CA*, and *RBMX*) were recurrently shared across patients, underscoring the predominantly patient-specific nature of subclonal evolution. Nevertheless, several genes disrupted by pathogenic or likely pathogenic subclonal variants converge on known drivers of luminal breast cancer biology and endocrine resistance^[7,50]^, including *PIK3CA, CDH1, GATA3, MAP3K1, SF3B1, TBX3*, and *TP53*. These findings align with our WES results, reinforcing their biological relevance in resistance pathways. Additional cancer drivers identified in subclones included chromatin remodelers (*CHD4, KMT2C, SMARCA2*), DNA damage response genes (*BRIP1, ATM, RTEL1*), and signaling molecules (*ABL1, BRAF, FGFR2*), reflecting multiple mechanisms through which clonal adaptation may proceed. Importantly, driver gene– stratified trajectories revealed that both intrinsic resistance (persistent baseline clones) and acquired resistance (variants emerging after treatment) are shaped by alterations in canonical cancer drivers as well as less-characterised genes, supporting the concept that AI resistance arises through both common and patient-specific routes. Notably, genes such as *DMD, ABCA13, CLSTN3*, and *PRDM1* were repeatedly annotated across clone categories, suggesting broader roles in tumour plasticity and potential roles as biomarkers, while ExpectoSc analyses implicated stromal, endothelial, breast, and immune compartments in mediating regulatory effects (**Figure 6, Supplementary Figure 16**). Our gene-centric results indicate treatment-specific selective pressures under sequential AI therapy. In luminal epithelium, *SMARCA1* was downregulated and *RBMX* strongly upregulated; persistence after letrozole involved baseline alterations in *ATR, PIK3CA, SF3B1*, whereas persistence after exemestane reflected emergent changes in *SF3B1* and *RBMX*. Differential behaviour of *SPEN/CHD4* (emerging on exemestane, lost on letrozole) and *FGFR2* (persisting on letrozole, eliminated by exemestane) highlights non-overlapping resistance routes. Conversely, clones bearing *FAT1, JAK1, ARID1B*, and *ATM* were eliminated, consistent with treatment sensitivity. Notably, ExpectoSc effect sizes were higher in clones persistent across both treatments or after exemestane, supporting functional impact and pointing to convergent pathways, PI3K signalling, chromatin remodelling, and splicing, as candidates for stratified therapeutic strategies.

Our pathway analyses revealed that resistance to aromatase inhibitors arises through a convergence of epigenetic, DNA damage, immune, and metabolic programs, while sensitivity is maintained through preserved checkpoint control and immune signaling. Pan-resistant clones, persisting under both letrozole and exemestane, demonstrated the most complex enrichment profile, spanning chromatin remodeling, histone modification, DNA repair, and EMT pathways. These were coupled with broad immune activation (interferon, JAK–STAT, TNF/NFκB, myeloid responses) and metabolic rewiring (fatty acid oxidation, glycolysis), as well as sustained hormone receptor signaling, including estrogen and androgen pathways. Such enrichment underscores the plasticity of resistant clones, integrating immune evasion, genomic instability, and steroid hormone signaling to maintain growth under endocrine pressure. Our observation of AR pathway involvement in resistant clones aligns with recent reports showing that exemestane and its metabolite directly interact with the androgen receptor, enhancing AR activation while suppressing ER signalling^[57]^. Beyond pathway-level enrichment, we also detect upregulation of *ITGAV* (Z-score 1.9–4.4) across all breast cell types in ExpectoSc, reinforcing AR-axis engagement at the gene level. This dual mechanism may explain why AR-positive tumours exhibit differential responses to steroidal versus non-steroidal AIs, and supports further investigation of AR as a determinant of treatment outcome and potential stratification marker in endocrine therapy.

Drug-specific resistance profiles revealed distinct mechanisms. Exemestane-only resistance was linked to telomere maintenance, RNA processing, oxidative stress, and acute inflammatory responses, while letrozole-only resistance showed a stronger metabolic and cell cycle signature but lacked immune activation, suggesting that suppression of immune pathways may facilitate exemestane effectiveness. Cross-resistance categories highlighted differential engagement of epigenetic reprogramming, MYC, and KRAS signaling, pointing to distinct but partially overlapping escape routes. In contrast, pan-sensitive clones preserved E2F- and TP53-driven cell cycle regulation, DNA repair, and immune signaling (IL2–STAT5, TNFα responses), suggesting that pathway silencing, rather than activation, underlies treatment failure.

From a clinical standpoint, enrichment of steroid hormone, PI3K/AKT/mTOR, and DNA repair pathways in resistant clones supports rational combinatorial strategies. We identified actionable alterations in PIK3CA (4 patients) and AKT1 (1 patient), with subclonal variants that persisted or re-emerged under sequential therapy, findings consistent with clinical benefit from PI3K inhibition in PIK3CA-mutated HR+/HER2− disease (SOLAR-1)^[58]^ and from AKT-pathway inhibition in PIK3CA/AKT1/PTEN-altered breast cancer^[59].^ Additional therapeutic opportunities were identified across PI3K/AKT/mTOR, CDK4/6^[60]^, DNA repair (*BRCA1/2, ATM*), and immune checkpoint pathways. These findings highlight that resistance to aromatase inhibition is driven by both convergent and drug-specific mechanisms, with actionable vulnerabilities in PI3K/AKT, chromatin remodeling, and immune pathways offering potential routes to overcome or prevent endocrine resistance.

A major strength of our study is the integration of bulk WES with subclonal reconstruction and validation using targeted scDNA-seq, allowing us to link genomic heterogeneity with treatment-specific trajectories in sequential AI therapy. The combined use of clinicogenomic associations, functional predictions, and pathway enrichment provided a multi-layered view of resistance biology, highlighting both convergent and patient-specific mechanisms. Nonetheless, the study is limited by the modest cohort size and the fact that subclonal reconstruction was feasible in only a subset of patients with sufficient tumour purity. Additionally, while our analyses identify candidate pathways and actionable targets, functional validation of these mechanisms remains necessary.

## Conclusion

This study provides novel insights into how clonal evolution shapes resistance to sequential AI therapy, identifying both shared drivers (*PIK3CA, SF3B1, RBMX*) and treatment-specific vulnerabilities. Our findings demonstrate that resistance emerges through a convergence of pathways, particularly PI3K/AKT/mTOR, chromatin remodelling, splicing, and immune signalling, while sensitivity is maintained through preserved checkpoint and DNA repair programs. The identification of actionable alterations in *PIK3CA* and *AKT1*, together with additional targets across DNA repair and immune pathways, highlights opportunities for rational combination strategies to overcome endocrine resistance. These results establish WES-guided subclonal inference as a feasible and clinically relevant approach, paving the way for more tailored therapeutic interventions in ER+ breast cancer.

## Acknowledgments

We acknowledge the contribution of Emma Rapp for the sub-analysing of samples of the Neoletexe trial using the Mission Bio Tapestri platform.

