## Supplementary Material for "Clonal selection supported by single cell Dna sequencing reveals hormonal adaptation and resistance in locally advanced breast cancer During Neoadjuvant Aromatase Inhibition"

### **Supplementary Tables**

#### **Supplementary Table 1.** Somatic mutation frequency obtained by exome sequencing in tumor biopsies obtained at baseline (T1), after 2 months (T2) and 4 months (T3) of aromatase inhibitor (AI) treatment. An asterisk (*) indicates that the number was averaged from two biopsies taken at the same time point. *EXE: exemestane; LET: letrozole; NA: data not available.*

| **Patient ID** | **T1** | **T2** | **T3** | **AI order** |
| --- | --- | --- | --- | --- |
| V23 | 183 | 292 | 304 | LET-EXE |
| J47 | 202 | 216 | 375* | EXE-LET |
| L73 | 1649 | 1101 | 1234 | LET-EXE |
| 053 | NA | NA | 245* | LET-EXE |
| Q88 | 331 | 88 | 241 | EXE-LET |
| L35 | 226 | 137 | 189 | LET-EXE |
| K12 | 149 | 150 | 208 | EXE-LET |
| M78 | 183 | 158 | 252 | LET-EXE |
| A66 | NA | NA | 196 | EXE-LET |
| A65 | 800 | 320 | 291 | LET-EXE |
| N29 | 612 | 367 | 730 | EXE-LET |
| L49 | NA | NA | 204 | LET-EXE |
| I37 | NA | NA | 242 | EXE-LET |
| E67 | 213 | 112 | 188* | LET-EXE |
| I89 | NA | 184 | 143 | LET-EXE |
| Z92 | NA | NA | 152 | EXE-LET |
| B32 | NA | NA | 293 | LET-EXE |
| W82 | NA | NA | 223 | EXE-LET |
| V27 | 1204 | 310 | 444 | LET-EXE |
| Z97 | NA | NA | 148 | LET-EXE |
| L11 | 263 | 598 | 315 | EXE-LET |
| T22 | 293 | 178 | 136 | LET-EXE |
| M98 | NA | NA | 285 | LET-EXE |
| W42 | 123 | 179 | 353 | LET-EXE |

#### **Supplementary Table 2**. Distribution of somatic mutations from 56 exomes into category and type. Indel: insertion or deletion; MNV: multi-nucleotide variant; SNV: single nucleotide variant.

|  | **Indel** | **MNV** | **SNV** | **Total** |
| --- | --- | --- | --- | --- |
| **Common** | 2 | 0 | 361 | 363 |
| **Benign** | 512 | 61 | 14,036 | 14,609 |
| **Likely Benign** | 32 | 7 | 1,087 | 1,126 |
| **Likely Pathogenic** | 16 | 4 | 291 | 311 |
| **Pathogenic** | 70 | 27 | 111 | 208 |
| **Unknown Significance** | 505 | 61 | 1,835 | 2,401 |
| Total | 1,137 | 160 | 17721 | 19,018 |
| MNV; Multi-Nucleotide Variant (substitution of >2 adjacent bases), SNV; Single Nucleotide Variant | | | | |

**Supplementary Table 3.** Summary of the number of variants and clones assessed and validated per patient. The number of timepoints analyzed, the number of total clones assigned by DPClust and the variants analysed by scDNA-sequencing are provided per patient.

| ID | N. of timepoints analysed with scDNA-seq | N. of DPClust clones | Variants in scDNA-seq panel |
| --- | --- | --- | --- |
| 2 | 1 | 6 | 16 |
| 3 | 3 | 9 | 141 |
| 8 | 1 | 5 | 24 |
| 9 | 2 | 1 | 11 |
| 11 | 2 | 5 | 25 |
| 12 | 2 | 9 | 67 |
| 15 | 3 | 6 | 55 |
| 17 | 2 | 3 | 13 |
| 21 | 3 | 7 | 77 |
| 24 | 3 | 4 | 31 |
| 29 | 2 | 5 | 21 |

N; number, scDNA-seq; single-cell DNA sequencing

### **Supplementary Figures**

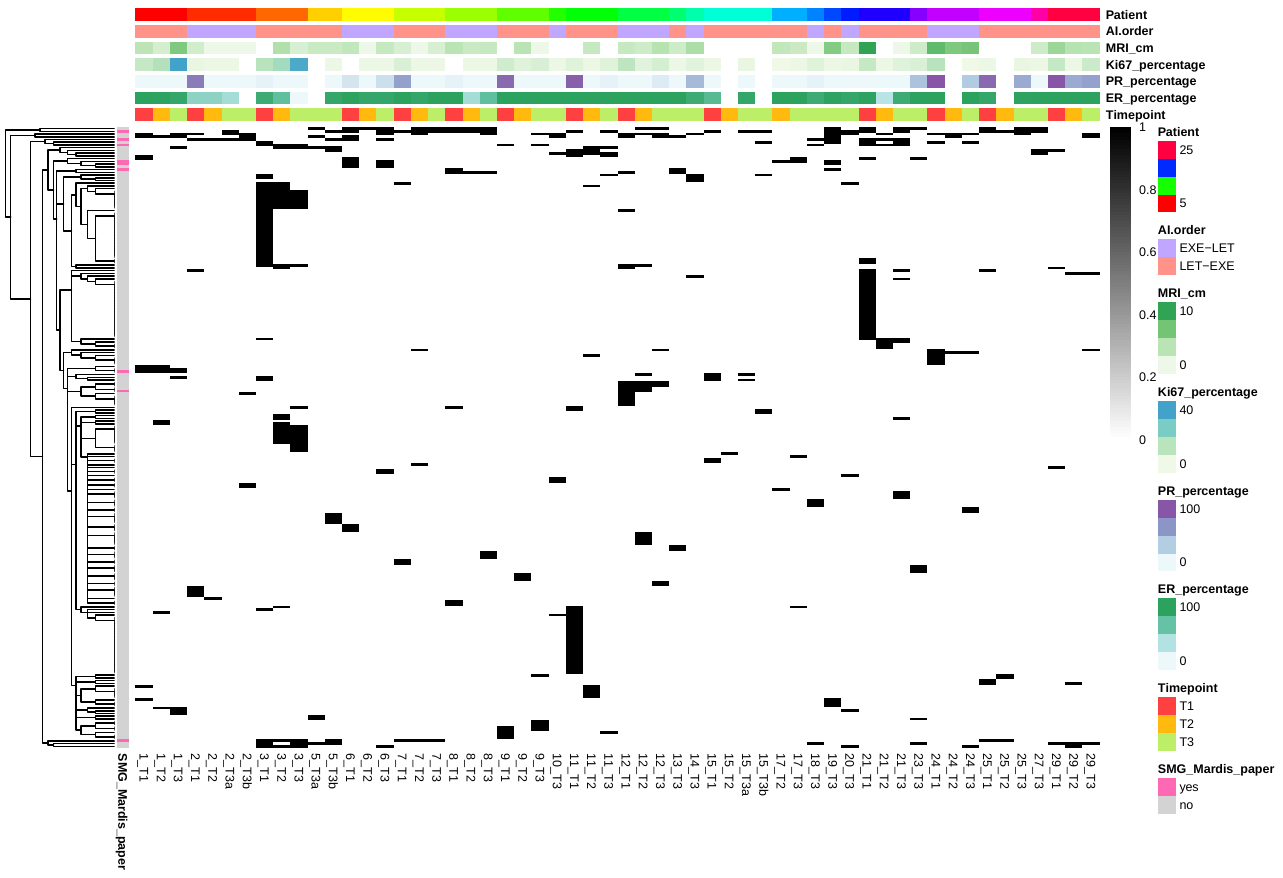
**Supplementary Figure 1.** Mutation matrix of all genes with somatic mutations classified as pathogenic or likely pathogenic (n=227 genes; rows) in the 56 exomes (columns). The columns are ordered according to patient number and time point. Black colour in the matrix indicates the presence of a mutation. White colour in the column annotation bar denotes missing data.

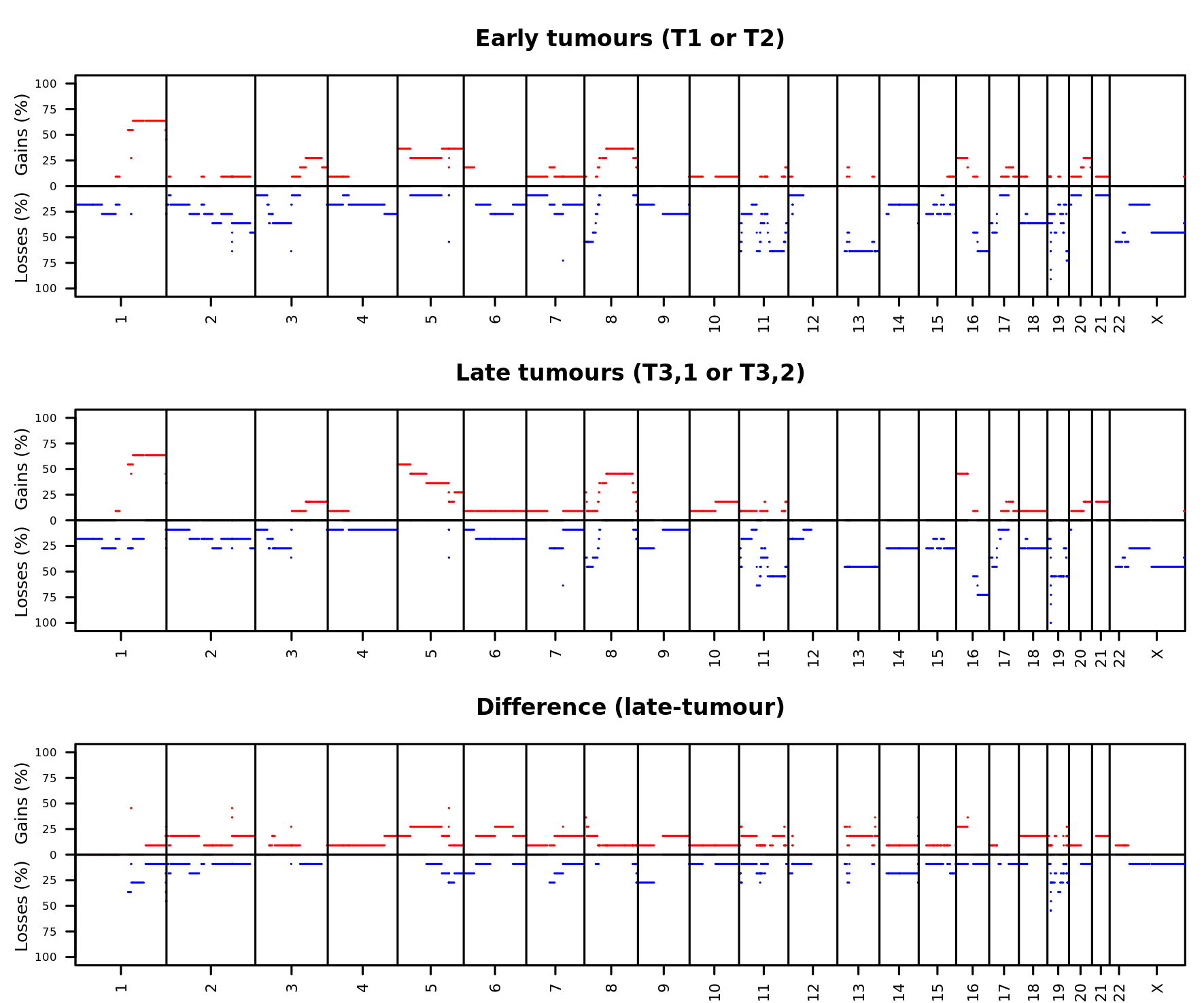

#### **Supplementary Figure 2**. Frequency of gains (red) and losses (blue) in early tumours (top panel) and late tumours (intermediate panel), as well as copy-number difference between early and late tumours (bottom panel).

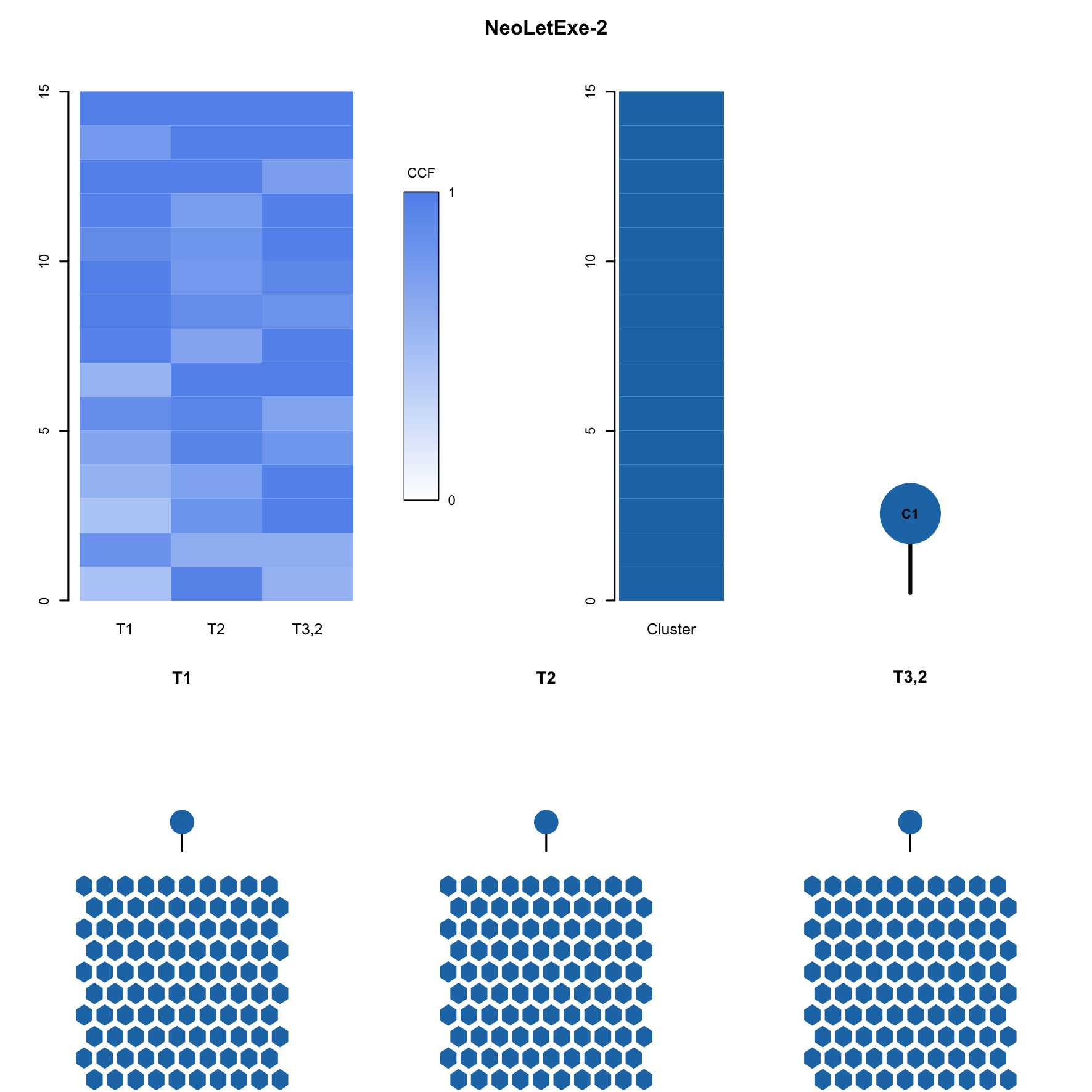

**Supplementary Figure 3**. **Subclonal composition and phylogenetic analysis of patient J47.** The top left panel shows cancer cell fraction (CCF) values for all of the SNVs (Y-axis) in all of the samples (X-axis). The top right panel shows, in relationship with the previous panel, mutation cluster(s) identified in each patient. The whole phylogenetic tree, showing relationships between different clones, are also displayed. The bottom panel shows, for each sample, which subclones are detected (in colours), as well as a representation of 100 cancer cells with nested colours, representing the presence of evolutionary sweeps.

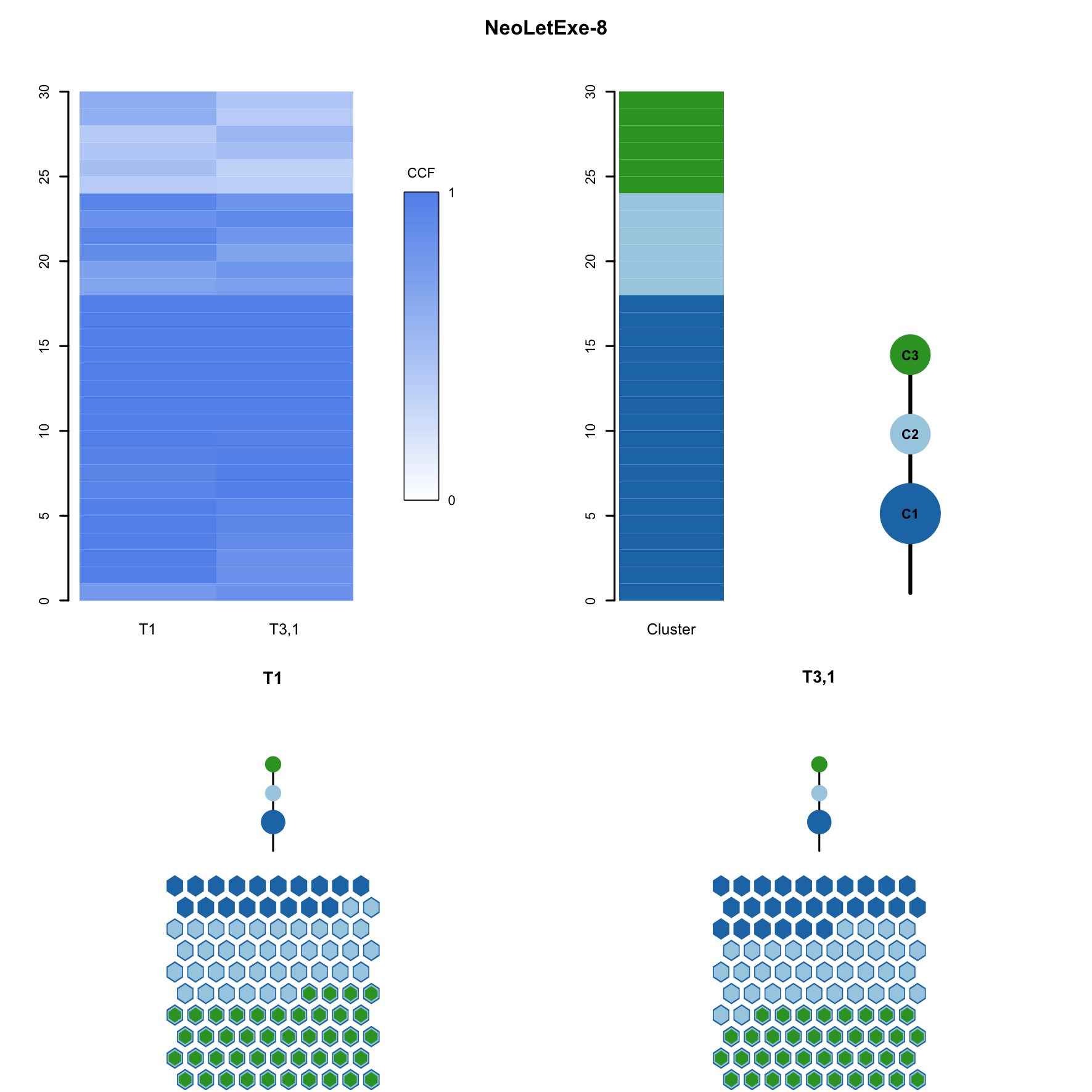
 **Supplementary Figure 4**. **Subclonal composition and phylogenetic analysis of patient K12.** The top left panel shows cancer cell fraction (CCF) values for all of the SNVs (Y-axis) in all of the samples (X-axis). The top right panel shows, in relationship with the previous panel, mutation cluster(s) identified in each patient. The whole phylogenetic tree, showing relationships between different clones, are also displayed. The bottom panel shows, for each sample, which subclones are detected (in colours), as well as a representation of 100 cancer cells with nested colours, representing the presence of evolutionary sweeps.

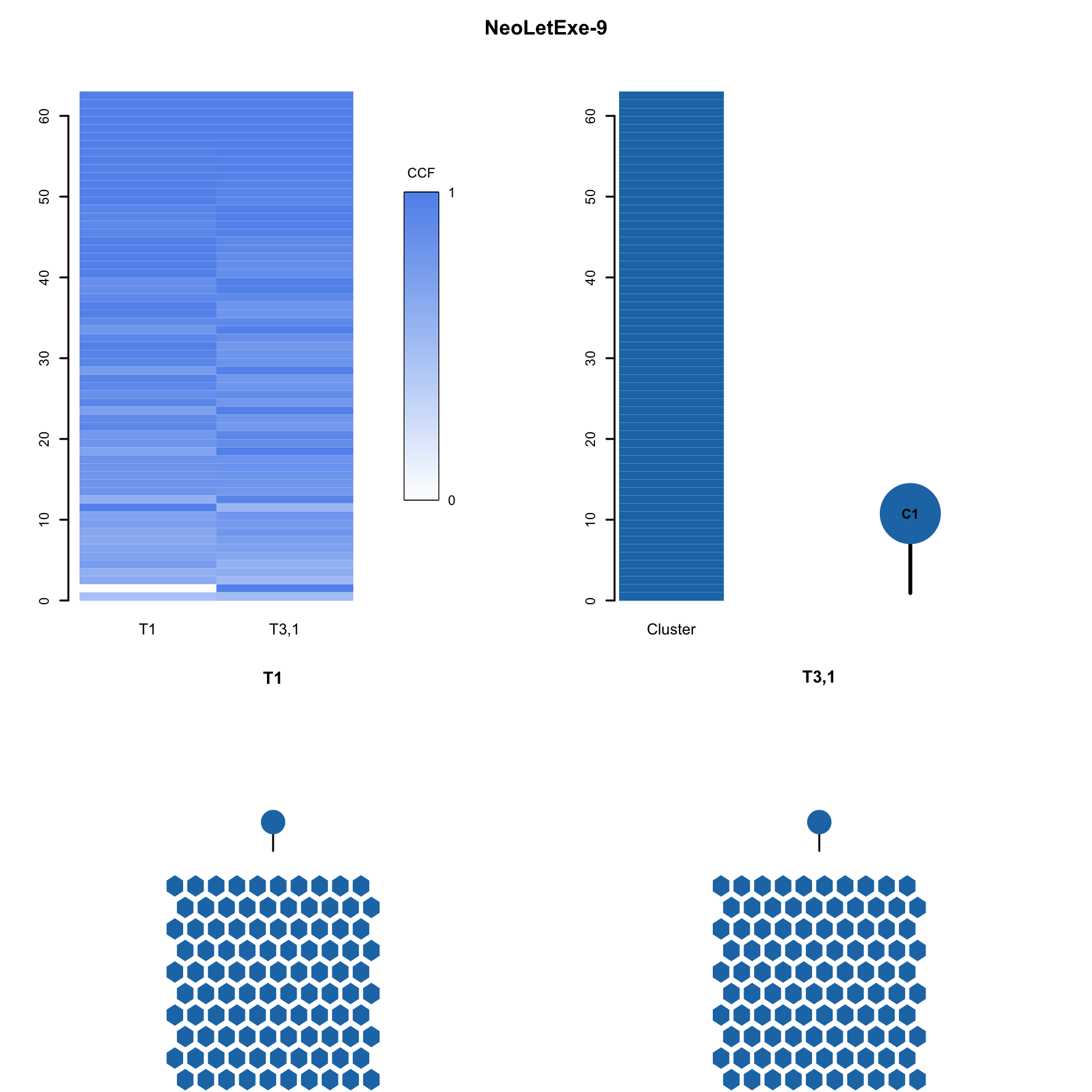
 **Supplementary Figure 5**. **Subclonal composition and phylogenetic analysis of patient M78.** The top left panel shows cancer cell fraction (CCF) values for all of the SNVs (Y-axis) in all of the samples (X-axis). The top right panel shows, in relationship with the previous panel, mutation cluster(s) identified in each patient. The whole phylogenetic tree, showing relationships between different clones, are also displayed. The bottom panel shows, for each sample, which subclones are detected (in colours), as well as a representation of 100 cancer cells with nested colours, representing the presence of evolutionary sweeps.

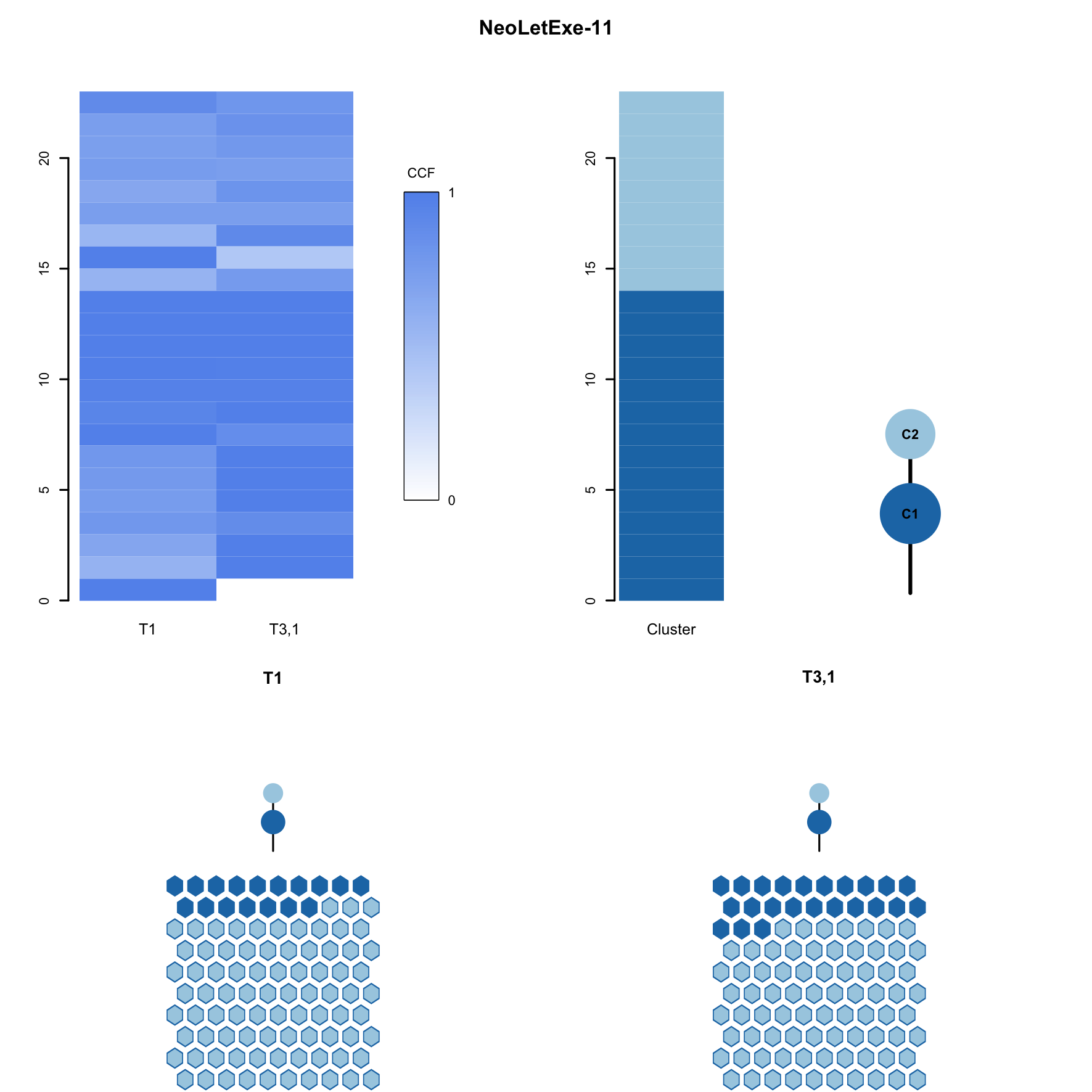
 **Supplementary Figure 6**. **Subclonal composition and phylogenetic analysis of patient A65.** The top left panel shows cancer cell fraction (CCF) values for all of the SNVs (Y-axis) in all of the samples (X-axis). The top right panel shows, in relationship with the previous panel, mutation cluster(s) identified in each patient. The whole phylogenetic tree, showing relationships between different clones, are also displayed. The bottom panel shows, for each sample, which subclones are detected (in colours), as well as a representation of 100 cancer cells with nested colours, representing the presence of evolutionary sweeps.

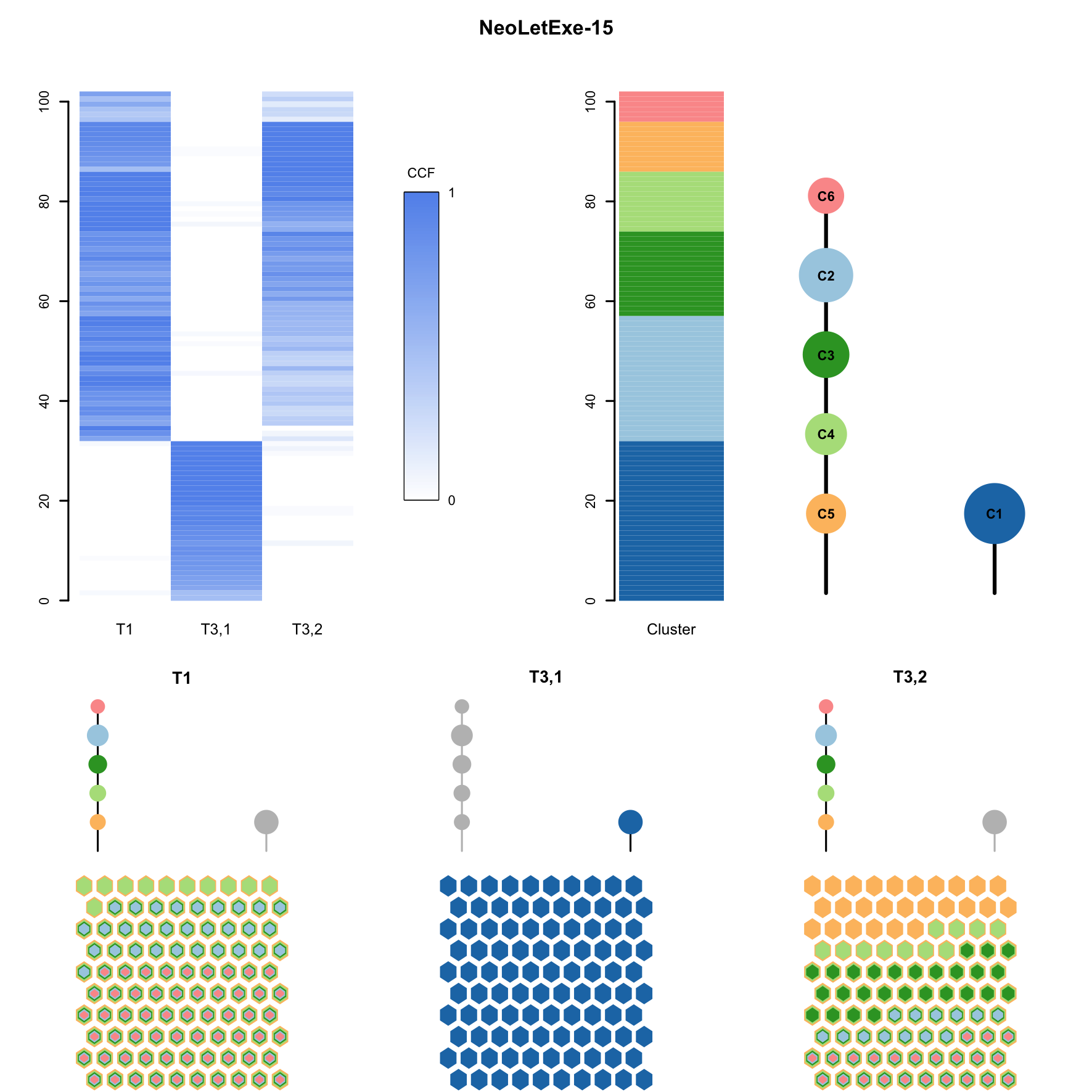

**Supplementary Figure 7**. **Subclonal composition and phylogenetic analysis of patient E67.** The top left panel shows cancer cell fraction (CCF) values for all of the SNVs (Y-axis) in all of the samples (X-axis). The top right panel shows, in relationship with the previous panel, mutation cluster(s) identified in each patient. The whole phylogenetic tree, showing relationships between different clones, are also displayed. The bottom panel shows, for each sample, which subclones are detected (in colours), as well as a representation of 100 cancer cells with nested colours, representing the presence of evolutionary sweeps.

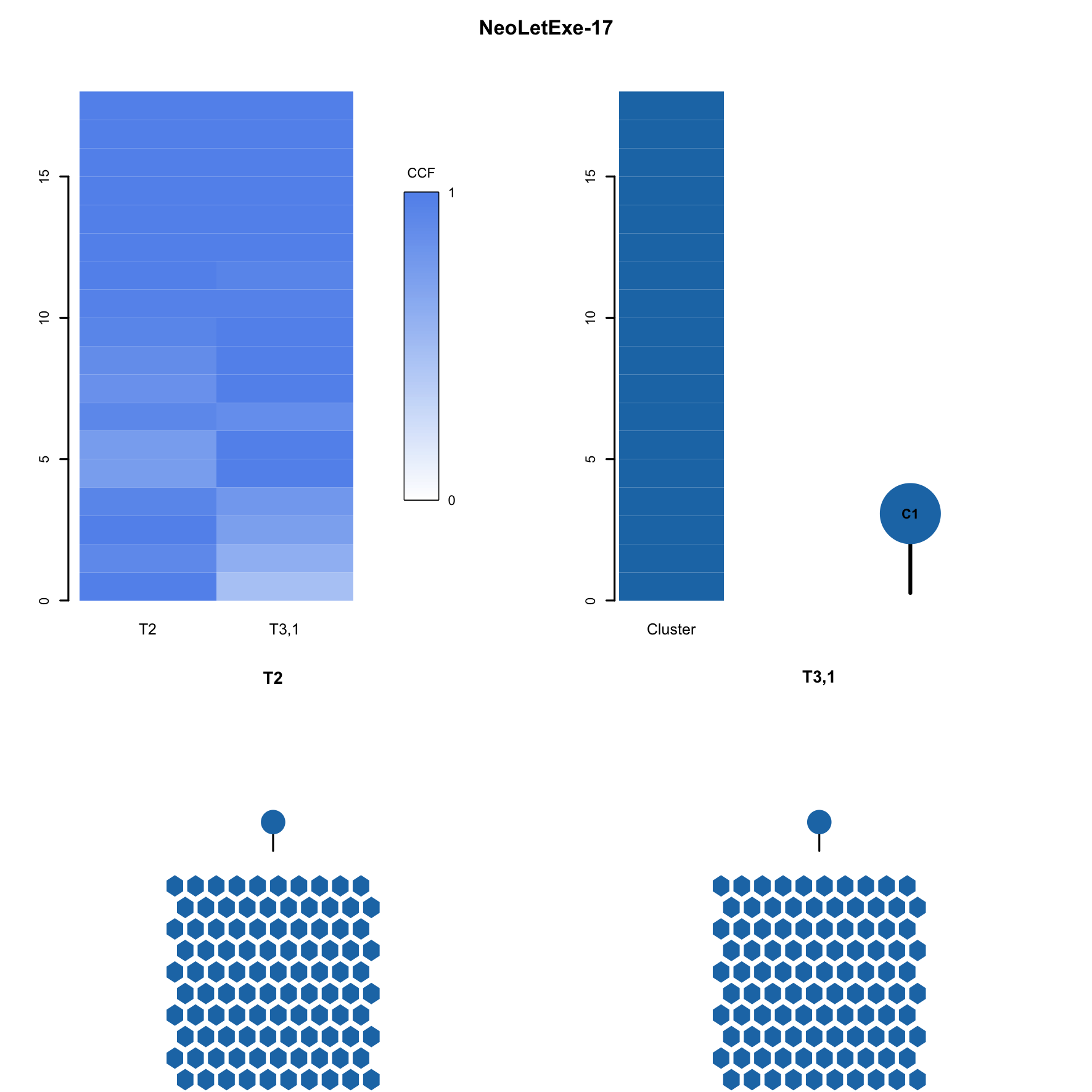

**Supplementary Figure 8**. **Subclonal composition and phylogenetic analysis of patient T70.** The top left panel shows cancer cell fraction (CCF) values for all of the SNVs (Y-axis) in all of the samples (X-axis). The top right panel shows, in relationship with the previous panel, mutation cluster(s) identified in each patient. The whole phylogenetic tree, showing relationships between different clones, are also displayed. The bottom panel shows, for each sample, which subclones are detected (in colours), as well as a representation of 100 cancer cells with nested colours, representing the presence of evolutionary sweeps.

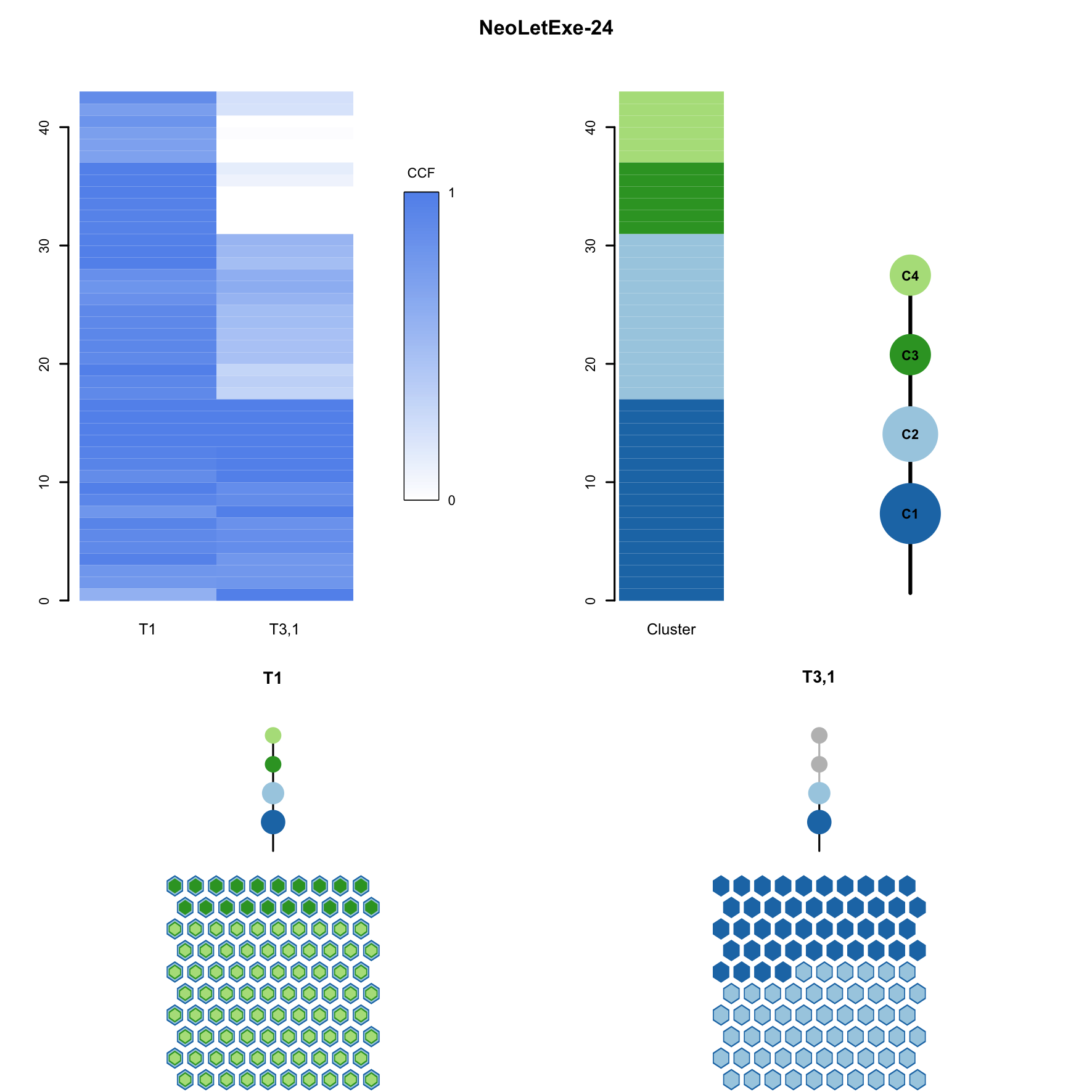
 **Supplementary Figure 9**. **Subclonal composition and phylogenetic analysis of patient L11.** The top left panel shows cancer cell fraction (CCF) values for all of the SNVs (Y-axis) in all of the samples (X-axis). The top right panel shows, in relationship with the previous panel, mutation cluster(s) identified in each patient. The whole phylogenetic tree, showing relationships between different clones, are also displayed. The bottom panel shows, for each sample, which subclones are detected (in colours), as well as a representation of 100 cancer cells with nested colours, representing the presence of evolutionary sweeps.

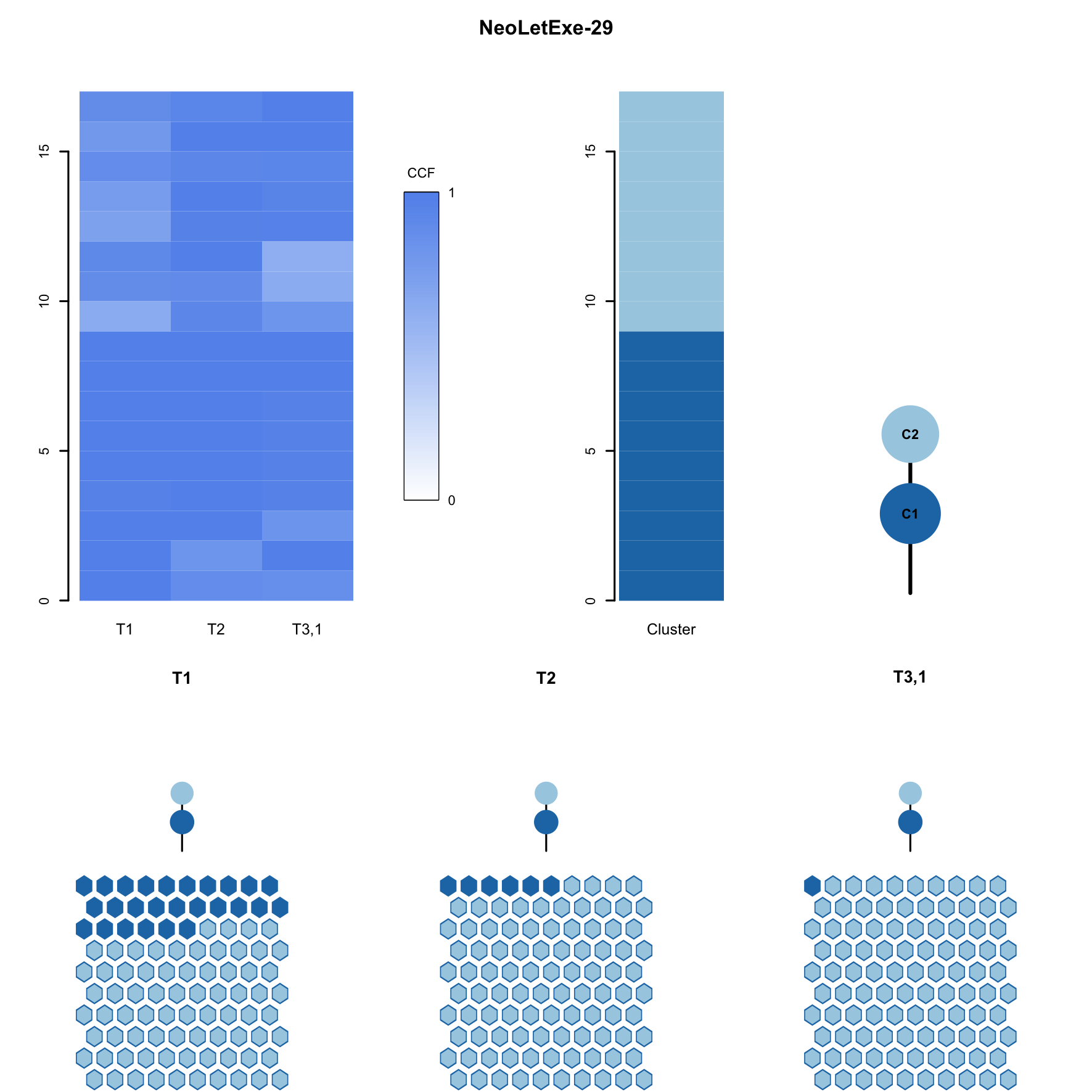

**Supplementary Figure 10**. **Subclonal composition and phylogenetic analysis of patient W42**. The top left panel shows cancer cell fraction (CCF) values for all of the SNVs (Y-axis) in all of the samples (X-axis). The top right panel shows, in relationship with the previous panel, mutation cluster(s) identified in each patient. The whole phylogenetic tree, showing relationships between different clones, are also displayed. The bottom panel shows, for each sample, which subclones are detected (in colours), as well as a representation of 100 cancer cells with nested colours, representing the presence of evolutionary sweeps.

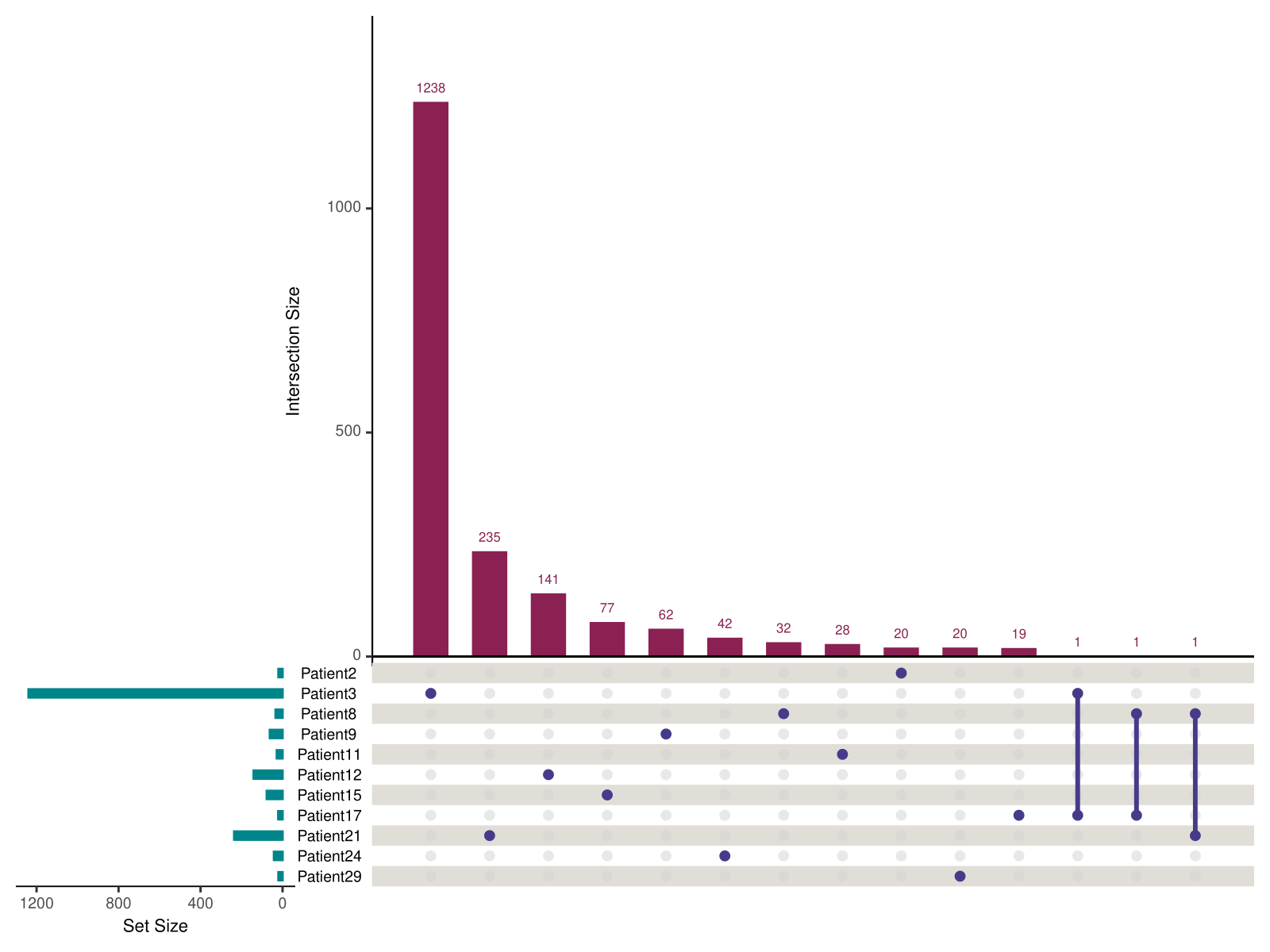

**Supplementary Figure 11.** **Co-occurrence of subclone variants between patients.** Interconnected dots represent shared entries, while single dots represent unique variants. The number of unique and intersection entries are represented by the pink bars, while the green bars on the left corner represent the total number of variants detected in all clones of each patient.

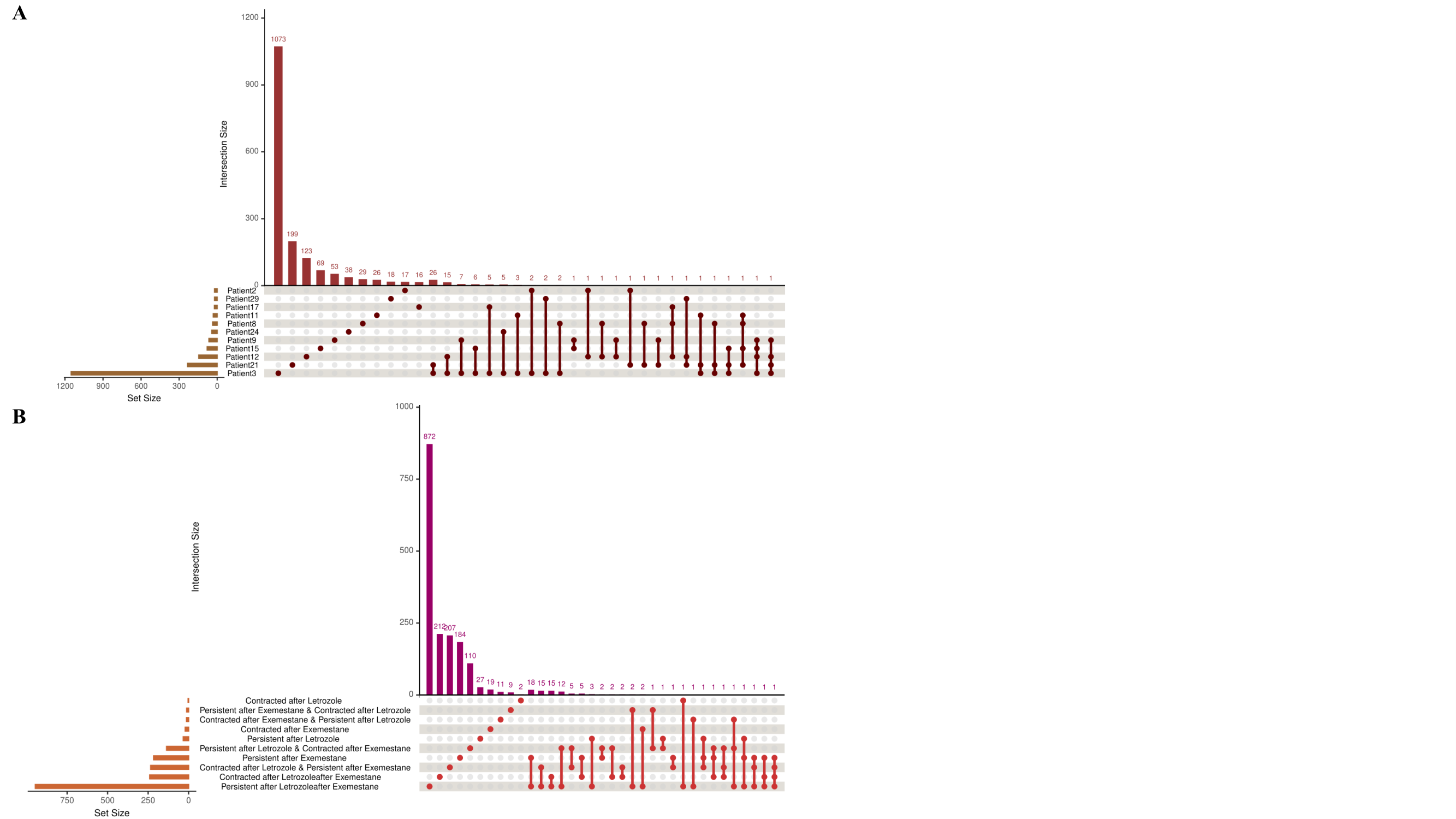

**Supplementary Figure 12.** **Co-occurrence of genes annotated to subclonal variants between patients and clone categories. (A)** Upset plot of genes annotated based on subclonal variants detected in patients. Interconnected dots represent shared entries; single dots represent unique variants. The number of unique and intersection entries are represented by orange bars. Yellow bars represent the total number of genes detected in all clones of each patient. **(B)** Upset plot of genes annotated in within the different clone categories.

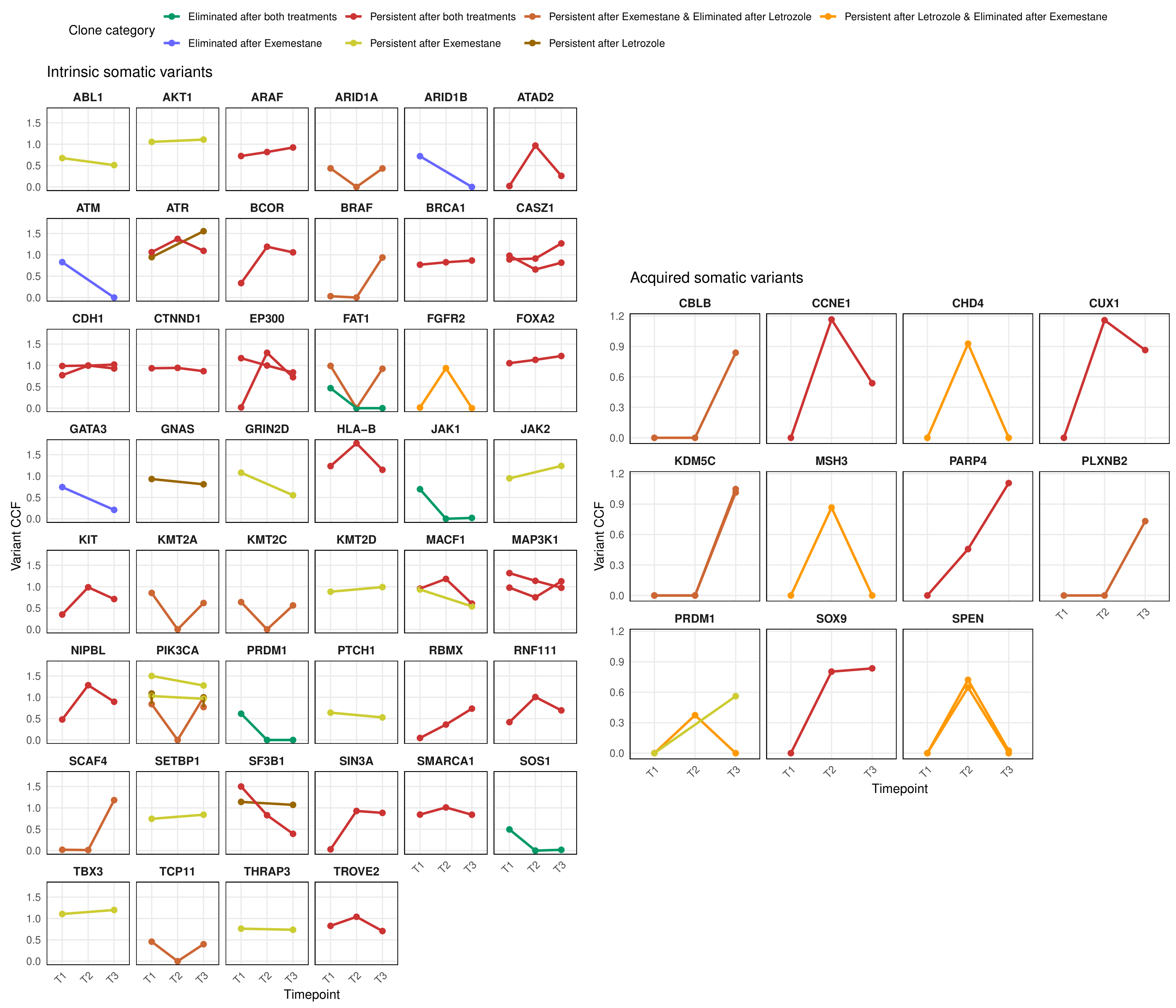

**Supplementary Figure 13.** **CCF trajectories of variants disrupting driver genes, across timepoints**. Dots represent clones measured in CCF (y-axis) across the timepoints, T1, T2 and T3 (x-axis). Clones are colour-coded according to the clonal category they belong to (see legend at the top).

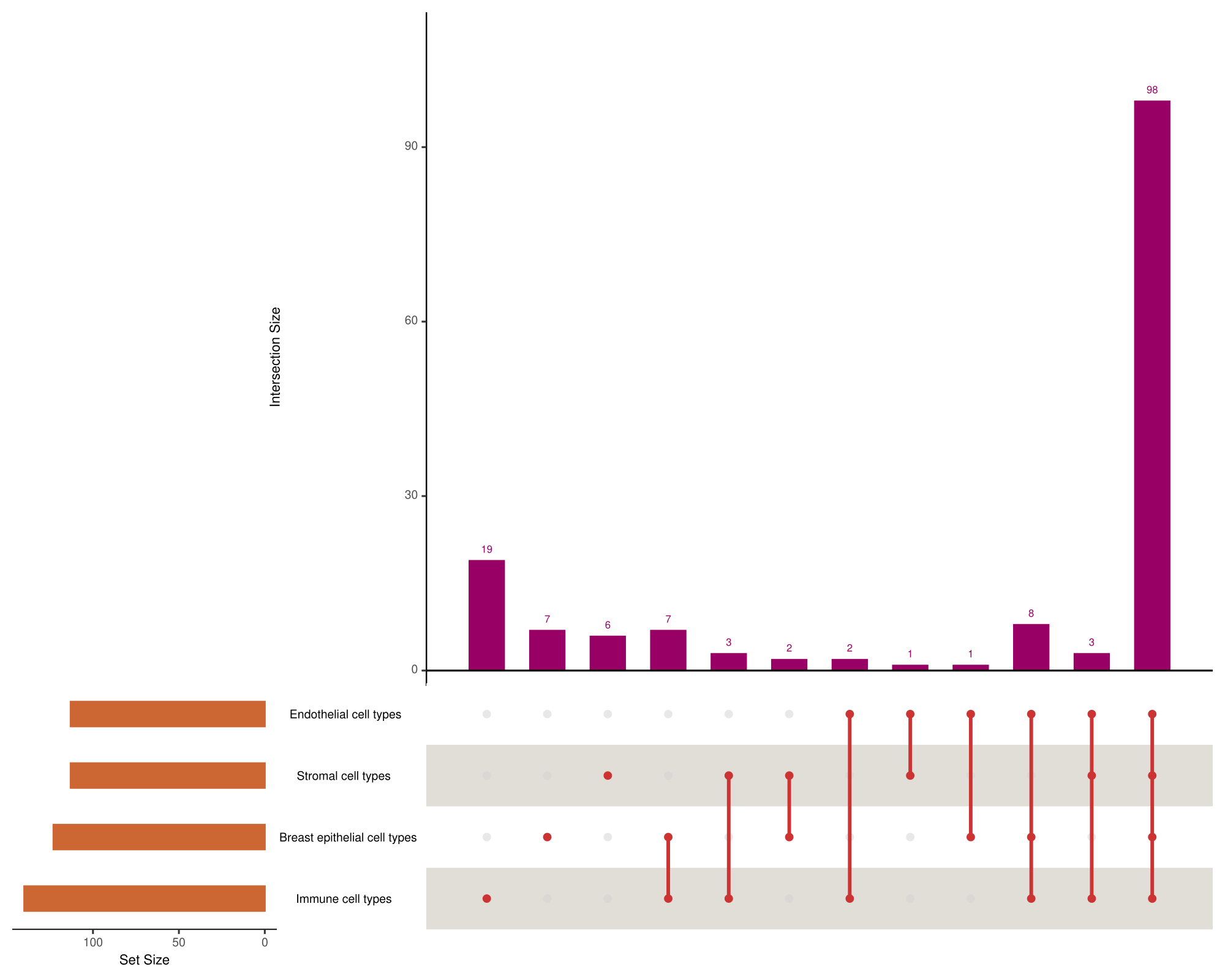

**Supplementary Figure 14**. **Co-occurrence of ExpectoSc predictions between cell types.** Upset plot showing the overlap of cell types (stromal, endothelial, breast, and immune) in which ExpectoSc QTL predictions are predicted to exert their regulatory effects.

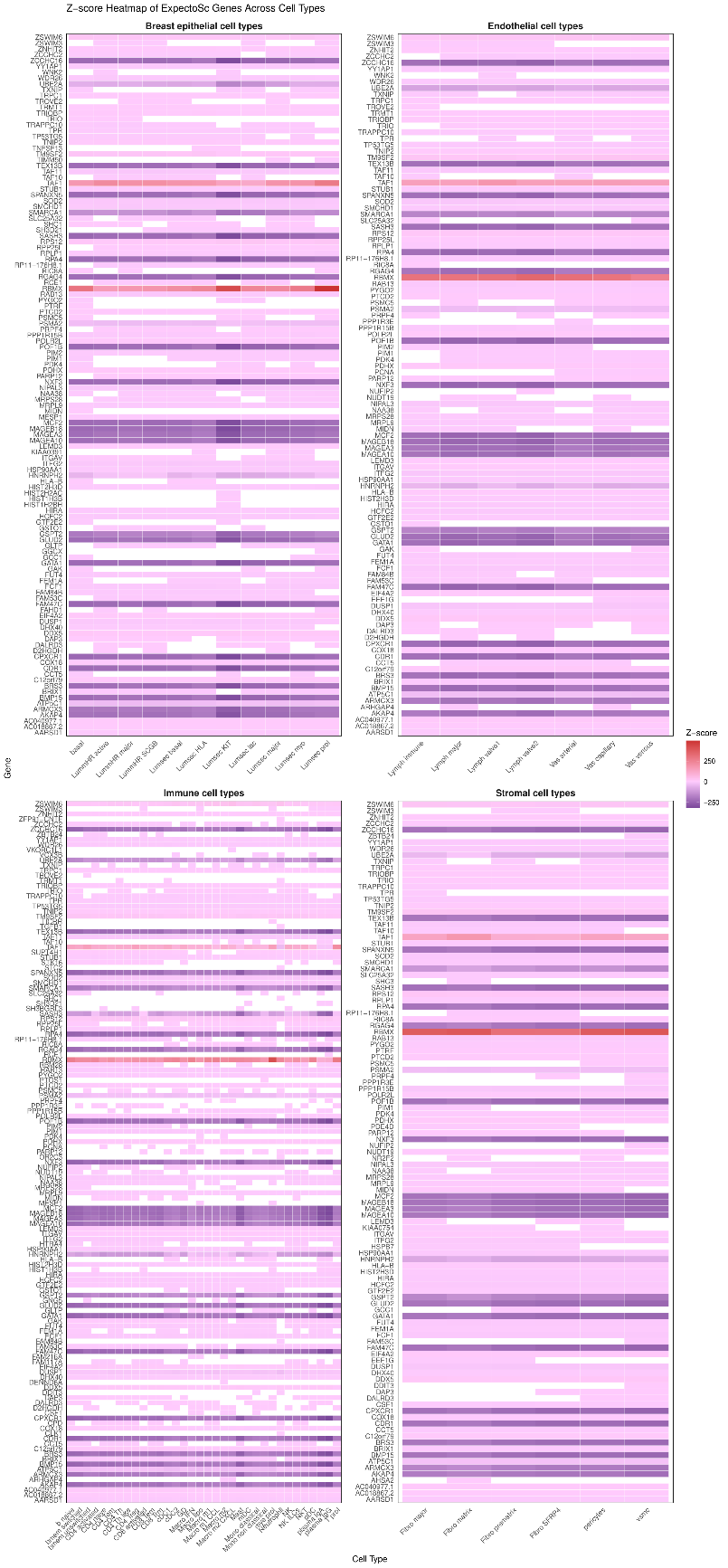

**Supplementary Figure 15.** **Heatmap of ExpectoSc predictions per tissue, measured in z-scores**. On the y-axis the genes impacted in expression by subclonal variant are shown, and on the x-axis the cell types corresponding to each tissue; Breast epithelial, Immune, Stromal and Endothelial cells, are provided.

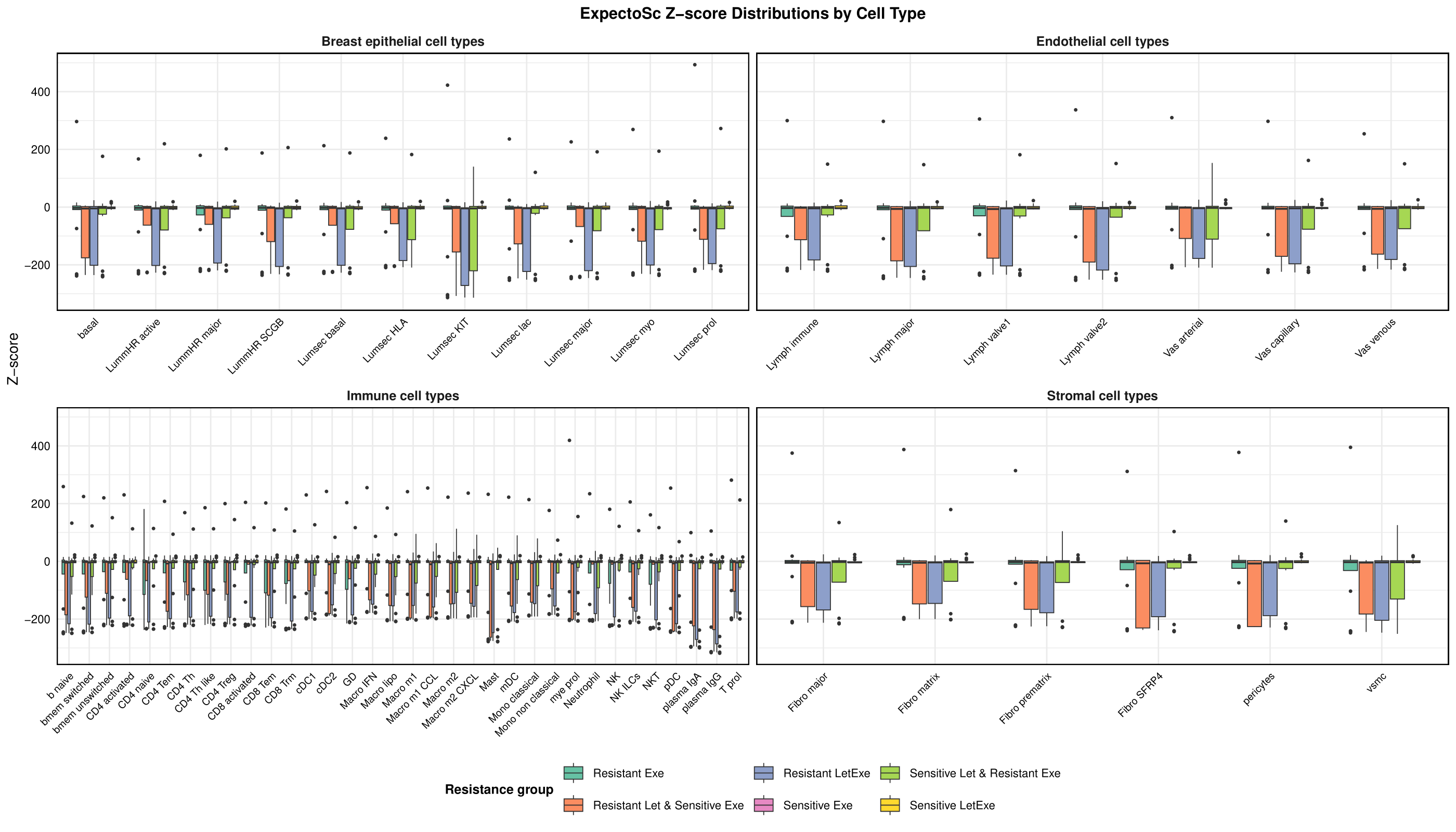
**Supplementary Figure 16**. **Distribution of ExpectoSc Z-scores predicting regulatory effects on gene expression, stratified by breast, endothelial, immune, and stromal** **cell types**. The y-axis shows the Z-score values, while the x-axis represents the specific cell types. Points are color-coded according to their associated resistant or sensitive clone categories.

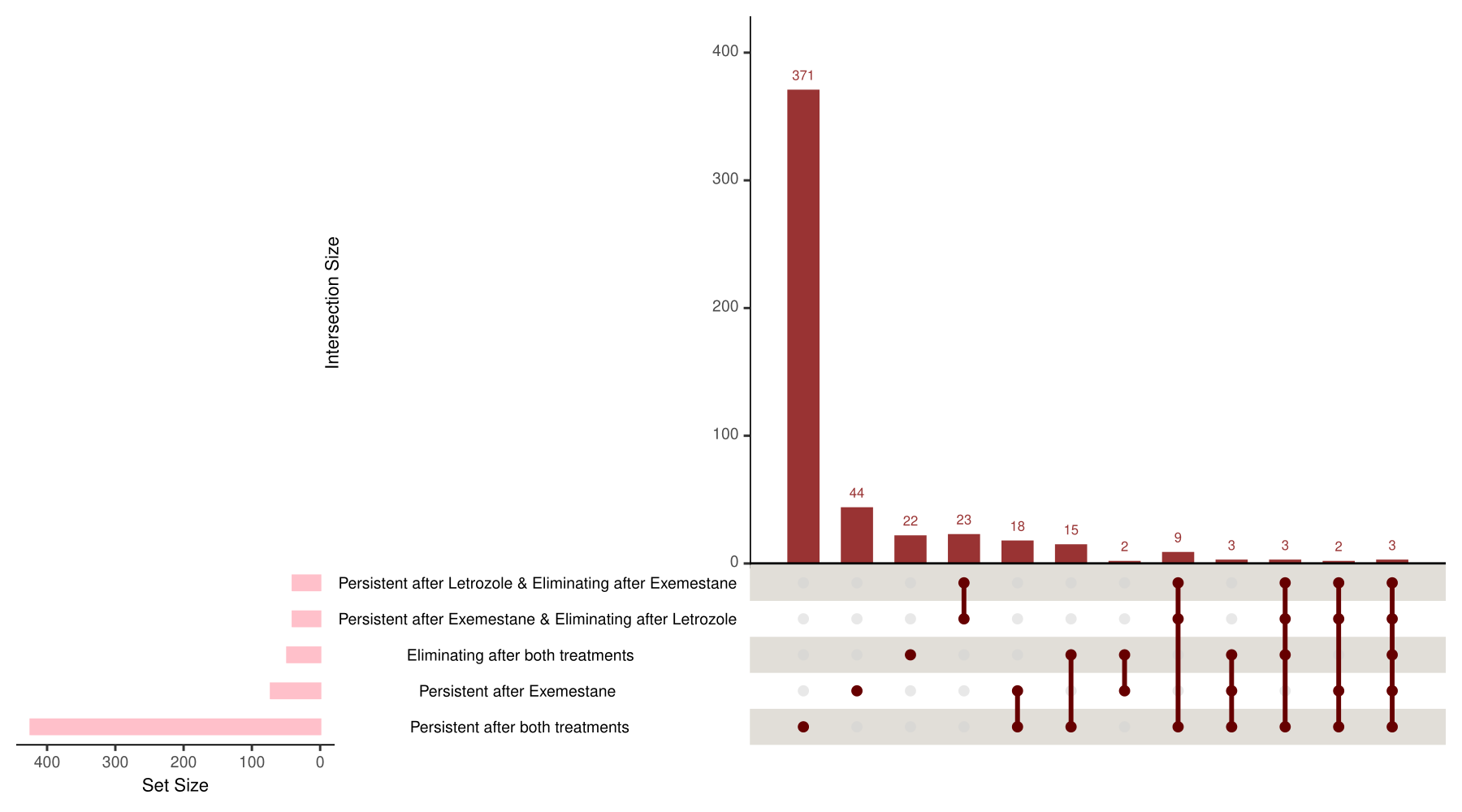

**Supplementary Figure 17.** **Co-occurrence of pathways found enriched at statistical significance, using Human Base between clone categories.** Interconnected dots represent shared entries; single dots represent unique variants. The number of unique and intersection entries are represented by orange bars. Pink bars represent the total number of genes detected in all clones of each patient.
